# Distinct intraspecific diversification dynamics in Neotropical montane versus lowland birds revealed by whole-genome comparative phylogeography

**DOI:** 10.1101/2024.11.01.620679

**Authors:** K. S. Wacker, B. M. Winger

## Abstract

Comparing phylogeographic patterns across different biogeographic regions can illuminate how different types of landscapes promote the formation of incipient species, providing insights into the evolutionary mechanisms underlying broadscale biodiversity gradients. The Neotropics are a global biodiversity hotspot, and the megadiverse Andes-Amazonia system has elevational gradients in both species richness and speciation rates. Here, we compare the phylogeographic dynamics of birds in the tropical Andes mountains versus the Amazonian lowlands using whole genomes from a congeneric set of colorful canopy frugivores (*Tangara* tanagers). We first show that Andean species have greater population structuring across their geographic ranges than Amazonian species. Next, we evaluated whether differences in geographic barrier efficacy and range size drive this elevated population-level divergence in the mountains. We find greater population differentiation and reduced gene flow across individual geographic barriers in the Andes. Furthermore, Andean species have consistently lower genetic diversity and smaller effective population sizes. Together, these results support a model of Neotropical diversification whereby incipient species form more readily in the mountains than the lowlands owing to more effective geographic barriers and smaller populations. These different phylogeographic processes across the Andes-Amazonia system carry implications for our understanding of the origin and maintenance of regional biodiversity gradients.

## Introduction

A major goal of evolutionary biology is to elucidate the microevolutionary mechanisms underlying macroevolutionary patterns (Rolland et al. 2023), including widespread biodiversity patterns like species richness and speciation rate gradients (Mittlebach et al. 2007, Schluter and Pennell 2017). Because the formation of new species typically requires a period of allopatry (Mayr 1942, Price 2008), understanding how levels of geographic population isolation and subsequent genetic divergence vary across the globe has the potential to reveal mechanisms underlying these patterns (Martin and McKay 2004, Eo et al. 2008, Smith et al. 2017). Historically, the study of incipient species formation across geographic barriers (i.e., comparative phylogeography) has focused on the response of a co-distributed assemblage to regional features of their shared landscape (Avise et al. 1987, Edwards et al. 2021, Provost et al. 2021); however, making these comparisons across multiple biogeographic regions is necessary to reveal how qualitatively different types of landscapes promote species formation. For example, contrasting mainland versus island taxa reveals differential effects of area and insularization on population differentiation and speciation between these types of landscapes (Mayr 1942, MacArthur and Wilson 1967, Diamond 1977, Loureiro et al. 2020, Naughton et al. 2024), improving our understanding of biodiversity patterns characterizing continents versus islands (Kier et al. 2009, Patton et al. 2021). Less work has been devoted to understanding how the microevolutionary processes of isolation, divergence, and speciation play out across different types of continental regions, such as mountains versus lowlands.

Within continental systems, in addition to latitudinal diversity gradients, there are well-characterized elevational biodiversity gradients. Specifically, species richness generally decreases with elevation globally (Graham et al. 2014), whereas speciation rates appear to increase with elevation across many branches of the tree of life (Esquerre et al. 2019, Testo et al. 2019, García-Rodríguez et al. 2021, Igea and Tanentzap 2021). These same patterns are particularly prominent in birds of the megadiverse Neotropical Andes-Amazonia system. This area holds more bird diversity than anywhere else in the world (Jenkins et al. 2013, Rahbek et al. 2019b) with species richness peaking in the Amazonian lowland and foothill regions, and decreasing with altitude (Terborgh et al. 1984, Kattan and Franco 2004, Quintero and Jetz 2018, Crouch et al. 2019). However, avian speciation rates have been reported to be lower in the Amazonian lowlands and increase with elevation in the Andes mountains (Weir 2006, Quintero and Jetz 2018). While there has been much study on comparative phylogeography within these mountain and lowland regions independently (Weir 2009, Smith et al. 2014, Winger 2017, Naka and Brumfield 2018), there are few studies assessing the microevolutionary dynamics of incipient species formation bridging the lowland-mountain gradient. Previously, Wacker and Winger (2024) demonstrated an elevational phylogeographic diversity gradient in Neotropical birds using mitochondrial DNA, whereby intraspecific population structure tends to be higher in Andean species than in Amazonian ones. While this suggests a greater propensity for population isolation in the Andes than in Amazonia, the mechanisms driving greater phylogeographic structure in the mountains remain untested. Clarifying the drivers of variation in phylogeographic divergence in Amazonia versus the Andes is important for understanding the origin and maintenance of biodiversity gradients in the most species-rich region in the world.

One fundamental reason phylogeographic divergence dynamics may differ across any two biogeographic regions is variation in the relative efficacy of regional barriers in promoting allopatry, with stronger and more stable barriers generating greater population divergence (Mayr 1942, Coyne and Orr 2004). For example, in Indo-Pacific island birds, wide oceanic barriers drive greater genetic differentiation than lowland terrestrial barriers (Pujolar et al. 2022). In the Andes-Amazonia system, extremely high levels of Andean topographic and climatic heterogeneity coupled with steep environmental gradients provide many opportunities for allopatric divergence (Lagomarsino et al. 2016, Rangel et al. 2018, Rahbek et al. 2019a). In particular, deep and arid valleys interrupting continuous humid forest tracts along a slope (Graves 1985, Killeen et al. 2007, Weir 2009, Winger and Bates 2015) and high elevation ridges separating different range slopes (Hazzi et al. 2018, Chesser et al. 2020), serve as physical barriers delimiting the ranges of montane species and populations. The isolating strength of these barriers is amplified by the narrow climatic niches characterizing tropical montane taxa (Janzen 1967, Cadena et al. 2012, Polato et al. 2018) and the linearity of Andean species’ geographic ranges, which presumably make it easier for a barrier to thoroughly fragment the range (Graves 1988). Despite the many factors yielding strong and stable geographic barriers in the mountains, diversification across the Andes has certainly involved episodes of dispersal across these barriers (Benham et al. 2015, Winger et al. 2015), especially when fluctuating climatic conditions cause increased connectivity of humid forest habitat, as during Pleistocene glacial cycles (Ramirez-Barahona and Eguiarte 2013, Hazzi et al. 2018).

By comparison, in lowland Amazonia, the Amazon River and its major tributaries act as physical barriers structuring species and populations (Wallace 1852, Ribas et al. 2012, Smith et al. 2014, Oliveira et al. 2017). Rearrangements of dynamic river networks via capture and avulsion events (Pärssinen et al. 1996, Almeida-Filho and Miranda 2007, Hayakawa and Rossetti 2015, Albert et al. 2018) can passively bring diverging populations back into geographic contact. These rearrangements occur over short timescales relative to the timeframe to speciation, and have been common in Amazonia since the establishment of the transcontinental Amazon River (Albert et al. 2018, Bicudo et al. 2019, Ruokolainen et al. 2019). Additionally, many species are capable of crossing rivers directly (Moore et al. 2008, Naka et al. 2022), potentially with the aid of river islands as dispersal stepping stones (Dornas et al. 2022). Populations isolated by wide lower river courses may in fact be connected near their narrower headwater regions (Weir et al. 2015, Moncrieff et al. 2024). Finally, like Andean environments, lowland Amazonia was also affected by Pleistocene climate cycling—glacial periods caused drying throughout the lowlands (Cheng et al. 2013, Wang et al. 2017) and species may have retracted into more stable, humid western Amazonian forests (Silva et al. 2019, Dalapicolla et al. 2024). Together, these features may render rivers semi-permeable or “leaky” to gene flow (Weir et al. 2024). Accordingly, we might expect that geographic barriers in the Andes mountains are relatively more effective at promoting population divergence than those in the Amazonian lowlands, although barrier efficacy across these biogeographic regions has not been directly assessed.

In addition to barrier-driven differences, consistent variation in population size may also affect population divergence within any two biogeographic regions. Effective population size (N_e_) determines the strength of genetic drift (Wright 1931, Charlesworth 2009), and under the nearly neutral theory smaller populations should have overall higher molecular substitution rates and diverge more readily from one another than larger populations (Ohta 1992, Bromham 2020). Effective population size is frequently approximated in empirical systems by levels of genetic diversity (θ=4N_e_μ) (Kimura and Crow 1964, Ellegren and Galtier 2016). Genetic diversity can vary among species owing to numerous biotic and abiotic factors, including life history strategies (Romiguier et al. 2014, Pegan et al. 2024), historical demographic fluctuations (Wright 1938), and the effects of linked selection (Leffler et al. 2012, Cutter and Payseur 2013). Importantly, geographic range size is also predicted to affect levels of genetic diversity via its influence on census population size (Leffler et al. 2012, Leroy et al. 2021). In the Andes-Amazonia system, the geographic area of the Andes mountains is substantially smaller than that of the Amazon basin. Consequently, montane taxa may have smaller population sizes and less genetic diversity than lowland taxa, which could cause a relative increase in neutral population divergence across barriers in the mountains.

How geographic barriers affect population differentiation is mediated by a species’ ecological and life history traits (Papadopoulou and Knowles 2016, Harvey et al. 2017a). Thus, comparing how qualitatively different landscapes affect phylogeographic dynamics can be most accurately accomplished when ecological differences are minimized across study taxa. This is particularly important for organismal traits affecting dispersal, as dispersal behaviors impact levels of population genetic differentiation through space and time (Palumbi 2003, Chua et al. 2017, Provost et al. 2021, Mascarenhas and Carnaval 2024). In this study, we evaluate phylogeographic divergence dynamics across a suite of eight *Tangara* tanager (Thraupidae) species which are widely distributed across mountain and lowland regions of the Neotropics. These colorful birds possess similar life histories and ecologies: all are small-bodied and highly social, regularly traveling in mixed-species foraging flocks to find fruit and arthropods in the canopy of humid forests (Isler and Isler 1999). Notably, diet and forest stratum lead to predictable differences in species’ genetic differentiation across landscapes, with canopy-dwelling frugivores exhibiting reduced population genetic divergence compared to understory insectivorous ones (Burney and Brumfield 2009, Miller et al. 2021). Birds in exposed canopy tend to demonstrate reduced neophobia and more exploratory behaviors than those living in the dark interior understory of forests, rendering them more likely to cross habitat gaps (Laurance 2004, Laurance et al. 2004, Lees and Peres 2009). Further, frugivorous species are more dispersive owing to their tracking of seasonally variable plant reproductive fruit resources (Greenberg 1981). Thus, we expect our canopy frugivore *Tangara* study species to have similar high dispersal potential, making them an ideal system for the evaluation of diversification dynamics in different biogeographic regions.

In this study, we characterize the phylogeographic divergence dynamics of Andean montane versus Amazonian lowland birds to evaluate the differential effects of well-known physical barriers and range size on the formation of incipient species across these two biogeographic regions. We use whole genomes sequenced at moderate coverage (∼6x) of 161 individuals across 8 species of *Tangara* tanagers with wide distributions throughout the Andes or Amazonia. Within this comparative phylogeographic framework, we test the following predictions. First, we predict that montane species will show relatively higher levels of population structuring than lowland species, owing to greater levels of topographic and climatic heterogeneity in the mountains (Wacker and Winger 2024). Second, we predict that montane species will have greater genomic differentiation and lower levels of dispersal across individual geographic barriers, supporting the hypothesis that barriers in the tropical mountains promote stronger allopatry than lowland river barriers. Lastly, we predict that Andean species will have less genetic diversity and smaller effective population sizes than Amazonian species, in accordance with their smaller range sizes. We discuss the implications of our results for the mechanistic basis of biodiversity gradients in this megadiverse Neotropical region.

## Materials and Methods

### Study system

We compared diversification dynamics between 4 Andean and 4 Amazonian species of *Tangara* tanagers (Passeriformes: Thraupidae). These congeneric canopy frugivores are closely related, sharing a common ancestor approximately 7.3 Ma (McTavish et al. 2024). The 4 Andean species (*T. nigroviridis*, N=18; *T. parzudakii*, N=19; *T. vassorii*, N=20; *T. xanthocephala*, N=20) are found at mid-elevation cloud forests, although *T. vassorii* has an expanded elevational distribution up to elfin forests near treeline. The 4 Amazonian species (*T. chilensis*, N=24; *T. mexicana*, N=21; *T. schrankii*, N=19; *T. velia*, N=20) are broadly distributed throughout humid lowland forests of the Amazonian lowlands, with the exception of *T. schrankii*, which is primarily found in western Amazonia (Billerman et al. 2022).

### Whole-genome sequencing libraries

We initially began with 166 samples from specimen-vouchered tissues housed in natural history collections (Supplementary Table S1). We isolated genomic DNA using a Qiagen DNeasy Blood and Tissue Kit. Individual whole-genome libraries were prepared based on a modified Illumina Nextera library preparation protocol (Schweizer et al. 2021). Samples were pooled and sequenced on two lanes of a NovaSeq X platform at the University of Michigan Advanced Genomics Core using paired-end 150 bp reads. This resulted in an average of 30.5 ± 7.7 million reads per sample (range 6.6 – 47.0 million). With a typical avian genome size of 1.1 Gb, this amounts to approximately 8.3x raw coverage.

Next, we cleaned and filtered raw reads in their fastq files. Using AdapterRemoval v2.3.1, we trimmed adapters, Ns, and low-quality bases from both ends using the options --trimns and –trimqualities (Schubert et al. 2016). Then, we performed poly-G trimming with fastp v0.23.2 and the --cut_right option for a sliding window approach (Chen et al. 2018). This step is necessary because two-channel sequencing systems like NovaSeq X call G bases when fluorescence signal is low, leading to an enrichment of poly-G tails (De-Kayne et al. 2021).

We aligned samples to a high-quality, annotated, chromosome-level reference genome of another species in the tanager family, *Camarhynchus parvulus* (GCA_901933205). We performed the alignment with BWA-MEM (Li 2013) and sorted reads with SAMtools (Danecek et al. 2021). We filtered aligned reads by first removing overlapping paired end reads with clipOverlap in bamUtil (Jun et al. 2015). Then, we used MarkDuplicates to tag duplicate reads and assigned all reads to new read-groups with AddOrReplaceReadGroups in picard. The resulting bam files were indexed and analyzed with flagstat in SAMtools. At this stage, we excluded samples with a mapping rate below 65% including duplicate reads from further analysis (N=5), leaving 161 samples in the final dataset.

Finally, we locally realigned these bam files around indels against the reference genome using GATK v3.7 (Van der Auwera et al. 2013). We created a list of potential indels for each species using RealignerTargetCreator, and then applied the GATK IndelRealigner tool to each sample. The resulting individual bam files were used in all further analyses. We calculated the final depth of coverage for each bam file by computing read depth at each high-quality filtered and aligned base with the SAMtools depth function.

### Mitochondrial genomes

Whole genome sequencing, even at low- or moderate-coverage, will yield full mitogenome sequences at high depth of coverage (Ye et al. 2014). We assembled full mitochondrial genomes from each individual sequencing library using NOVOplasty v4.3.4 (Dierckxsens et al. 2017), using a single ND2 gene sequence from a conspecific on GenBank as the seed (Supplementary Table S2). Of the 161 samples passing nuclear filters, 159 mitogenomes assembled correctly (*T. vassorii* LSUMNS B-593 and *T. velia* FMNH 389278 could not be assembled by NOVOplasty). Mean mitogenome coverage was 7705 ± 5913 copies (range 49 – 30,428 copies); species-level assemblage summaries are given in Supplementary Table S2. From these contigs, we extracted all 13 protein-coding mitochondrial genes using Geneious v2021.2.2.

Mitochondrial gene sets were used for two purposes: data quality control and population structure analyses. First, we BLASTed the ND2 gene for each sample, verifying species identity and ruling out potential specimen mis-identification or library contamination. We also visually inspected gene alignments with DECIPHER in R (Wright 2016) to rule out the presence of chimeras or hybrids. All mitogenomes passed this quality control check, and we concatenated genes into a complete 13 protein-coding gene set alignment for each species.

Second, we created phylogenetic trees to assess levels of intraspecific population structure. Most Neotropical phylogeographic studies to-date have used mitochondrial loci, and we wanted to contextualize levels of structuring in *Tangara* within this body of work. We created ultrametric ND2 gene trees from in BEAST 2.6.6 (Bouckaert et al. 2019), and assessed discrete structure in the ND2 trees with GMYC following (Wacker and Winger 2024). Then, we created full mitogenome phylogenies with the 13 protein-coding gene set alignment. We partitioned alignments by gene and codon, and constructed ML phylogenies in IQ-TREE 2 with automatic model detection (MFP+MERGE) and 1000 ultrafast bootstrap replicates (Minh et al. 2020).

### Nuclear genome datasets

We conducted all analyses of nuclear genomes in a genotype-likelihood framework to account for sequencing errors and base-calling uncertainties (Nielsen et al. 2011). We created two datasets for analyses of the nuclear genomes: one containing SNPs and one based on all autosomal sites (variant and invariant). We called SNPs in ANGSD v0.941 (Korneliussen et al. 2014) using the SAMtools genotype likelihood model (-GL 1), at the 0.05 significance level with a minimum MAF of 0.05. We filtered the resulting beagle files with ngsParalog v1.3.2 to remove SNPs with a high likelihood of occurring in duplicated or mis-mapped regions (Linderoth 2018). The number of filtered SNPs per species is listed in Table 1.

**Table 1.** Summary of study species and whole-genome sequencing statistics. Total reads = average total number of paired-end reads per sample. Mapping rate = average percentage of reads per sample mapping to the *Camarhynchus parvulus* reference genome. Filtered site depth = average per-base site depth of all sites in the bam file of each sample used in subsequent analyses. Filtered SNPs = number of SNPs for each species inferred by ANGSD at the 0.05 significance threshold that passed further filtering with ngsPara. Subsampled genome sites = the number of autosomal sites (variant and invariant) used for all analyses requiring the site frequency spectrum in ANGSD.

| Species | Region | N | Total reads | Mapping rate | Filtered site depth | Filtered SNPs | Subsampled genome sites |
| --- | --- | --- | --- | --- | --- | --- | --- |
| *T. chilensis* | Amazonia | 24 | 55,594,919 ± 16,321,847 | 84.3% ± 4.8% | 4.95 ± 1.47 | 69,780,156 | 111,205,357 |
| *T. mexicana* | Amazonia | 21 | 53,958,149 ± 12,849,476 | 86.5% ± 3.4% | 4.98 ± 1.13 | 35,989,731 | 111,231,300 |
| *T. nigroviridis* | Andes | 18 | 71,790,809 ± 13,049,380 | 89.8% ± 2.5% | 6.79 ± 1.18 | 29,553,375 | 111,301,155 |
| *T. parzudakii* | Andes | 19 | 59,681,121 ± 10,667,571 | 87.7% ± 2.3% | 5.56 ± 0.98 | 26,793,071 | 111,239,831 |
| *T. schrankii* | Amazonia | 19 | 68,501,805 ± 14,575,009 | 89.0% ± 3.6% | 6.46 ± 1.48 | 54,020,062 | 110,143,042 |
| *T. vassorii* | Andes | 20 | 56,618,951 ± 15,146,929 | 86.7% ± 3.4% | 5.27 ± 1.27 | 29,657,098 | 111,391,048 |
| *T. velia* | Amazonia | 20 | 63,189,998 ± 13,174,941 | 83.9% ± 14.2% | 5.38 ± 1.48 | 34,992,015 | 111,401,137 |
| *T. xanthocephala* | Andes | 20 | 60,781,970 ± 18,534,379 | 87.1% ± 5.3% | 5.62 ± 1.62 | 19,283,036 | 111,385,456 |

We also created a dataset of all sites (variant and invariant) within individual libraries per species, following best practice for population genetic inference from site frequency spectra (Nielsen et al. 2012, Korunes and Samuk 2021). Due to computational constraints when analyzing whole genome data in a genotype likelihood framework, these datasets comprised a random subsample of approximately 10% of the entire genome for tractability. We subsampled genomes using scripts modified from https://github.com/markravinet/genome_sampler, selecting random windows of 2kb that are located at least 10kb apart. We removed sites flagged as being paralogous or mismapped with ngsParalog, as well as sites mapping to the Z chromosome, because sex chromosomes have different effective population sizes and expectations for genetic diversity than autosomes. The resulting BED files were subsequently applied as site filters in ANGSD to restrict analyses to this subset of autosomal DNA.

### Evaluating levels of intraspecific population structure

We used the SNP dataset to evaluate population structure in three ways. First, we ran principal component analyses with PCAngsd v1.10 (Meisner and Albrechtsen 2018) to visually assess how individuals cluster in PC space relative to their geographic population (see *Evaluating strength of barriers* and Figure 1 for descriptions of geographic populations). Second, we determined the amount of discrete intraspecific structure by inferring the number of ancestry clusters (K) in each species. We ran admixture analyses with the --admix call in PCAngsd, which automatically infers the best-fitting K. We ran additional admix analyses manually setting K=2-5 to visually compare different numbers of ancestry clusters. We also assessed discrete structure with K-means clustering using DAPC with the *find.clusters()* function in adgenet (Jombart et al. 2010). Third, we assessed the strength of continuous isolation by distance (IBD) in each species using the pairwise genetic covariance matrix generated in PCAngsd. We calculated the geographic distance between all pairs of individuals within each species using geosphere in R (Hijmans 2016). We regressed pairwise values of 1 – genetic covariance against geographic distances (Novembre and Stephens 2008), and used the slope of the linear regression as the strength of isolation by distance, β_IBD_ (Singhal et al. 2018, Pegan et al. 2024). We further calculated Mantel statistics as another measure of the correlation between genetic distance and geographic distance for each species using the *mantel()* function in vegan (Oksanen et al. 2024).

**Figure 1.**
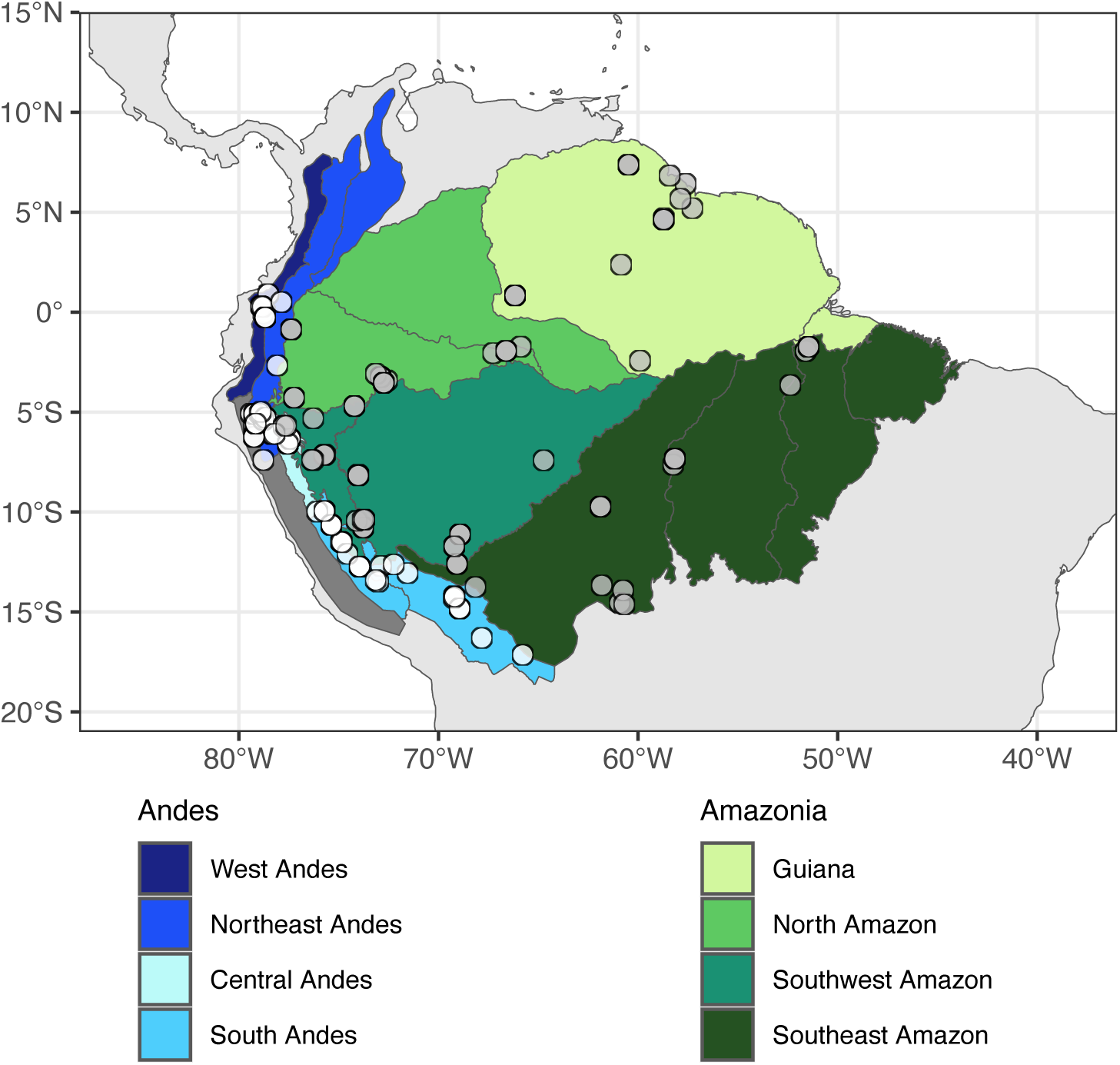
Map of study system. Amazonian samples (N=84) are represented by gray circles, and Andean samples (N=77) are represented by white circles. Geographic populations used to assess barrier efficacy are distinguished by various shades of green (Amazonian lowlands) and blue (Andean mountains). Specifically, in the mountains, samples in the West Andes are isolated on their slope from samples in the Northeast Andes; the Northeast Andes population is separated by the Marañón Valley barrier from the Central Andes in Peru; and the “Central Andes” (operationally defined for this study) are separated by the Huallaga Valley barrier from the South Andes population. The dark gray region of the Andes is the unsampled and dry west Peruvian slope. In the lowlands, the Guiana population is separated from the North Amazon population to the west by the Negro River barrier; the North Amazon population is separated from the Southwest Amazon population to the south by the Upper Amazon River barrier; the Southwest Amazon population is separated from the Southeast Amazon population by the Madeira River barrier; and the Southeast population (representing a grouping of interfluves) is separated from the Guiana population by the Lower Amazon River barrier.

### Evaluating strength of barriers

In order to assess how well barriers within each biogeographic region promote isolation, we evaluated pairwise population differentiation (F_ST_) and estimates of gene flow from demographic modeling across individual landscape barriers. This required *a priori* designations of geographic populations for our samples. We defined geographic populations as those separated by physical barriers with well-characterized and strong roles in structuring populations or species assemblages (Da Silva et al. 2005, Weir 2009, Hazzi et al. 2018), and included populations in analyses for a given species only if we had at least 2 samples within that region for that species. Geographic populations, the barriers separating them, and population sample sizes are summarized in Figure 1 and Table 2. This principal set of geographic population definitions was constructed to maximize sample size within each population while assessing the efficacy of major regional barriers, but required us to bin some individuals across other landscape features (mountain valleys or rivers) that could also function as barriers. To evaluate the sensitivity of our results to population groupings, we ran multiple alternative population scenarios. In the Andes, we bisected the “Southern Andes” population into two populations across the Apurímac Valley to evaluate this valley as an additional barrier (Supplementary Figure 8). In the lowlands, we created two alternative scenarios affecting western Amazonia: first, we changed the boundary of the Upper Amazon course from the Marañón River mainstem to the Ucayali River headstream, and second, we isolated samples between the Marañón and Ucayali as a separate fifth (“Huallaga”) lowland population (Supplementary Figure 9).

**Table 2.** Summary of geographic populations and barriers analyzed in this study. The strengths of individual geographic barriers (“Barrier”) were analyzed by pooling samples into predefined geographic populations (“Population 1” and “Population 2”). The number of samples per species across a particular barrier is also given, with sample size for Population 1 on the left side of the slash and sample size for Population 2 on the right side of the slash.

| Amazonia |  |  |  |  |  |  |
| --- | --- | --- | --- | --- | --- | --- |
| Population 1 | Population 2 | Barrier | *T. chilensis* | *T. mexicana* | *T. schrankii* | *T. velia* |
| Guiana | North Amazon | Negro River | 3 \| 6 | 5 \| 3 | -- | -- |
| North Amazon | Southwest Amazon | Upper Amazon | 6 \| 11 | 3 \| 7 | 5 \| 11 | -- |
| Southwest Amazon | Southeast Amazon | Madeira River | 11 \| 4 | 7 \| 6 | 11 \| 2 | 10 \| 3 |
| Southeast Amazon | Guiana | Lower Amazon | 4 \| 3 | 6 \| 5 | -- | 3 \| 6 |
| Andes |  |  |  |  |  |  |
| Population 1 | Population 2 | Barrier | *T. nigroviridis* | *T. parzudakii* | *T. vassorii* | *T. xanthocephala* |
| West Andes | Northeast Andes | Slope | 3 \| 5 | 5 \| 5 | 2 \| 6 | -- |
| Northeast Andes | Central Andes | Marañón Valley | 5 \| 4 | 5 \| 4 | 6 \| 2 | 8 \| 4 |
| Central Andes | Southern Andes | Huallaga Valley | 4 \| 6 | 4 \| 5 | 2 \| 10 | 4 \| 8 |

To evaluate F_ST_ across these barriers, we created site allele frequency spectra in ANGSD for each geographic population based on all autosomal sites (variant and invariant) with the -doSaf command. Next, we generated folded two-dimensional site frequency spectra (2D-SFS) for pairs of populations across particular barriers using winsfs (Rasmussen et al. 2022). We then used the 2D-SFS to calculate genome-wide F_ST_ using realSFS in ANGSD with the call -whichFst 1 (i.e., the Bhatia estimator, which is preferred for small sample sizes; Bhatia et al. 2013). We calculated F_ST_ for all alternative geographic population scenarios (2 Andean and 3 Amazonian) and compared mountain versus lowland differentiation for all 6 possible regional comparisons; results are unaffected by the choice of population designations (Supplementary Figure 10), and we present only the principal set of populations here (Figures 4-5).

Next, we compared levels of gene flow across particular barriers with demographic modeling in the program GADMA v2.0 (Noskova et al. 2023), employing moments as the engine for demographic inference (Jouganous et al. 2017). For each pair of populations separated by a barrier, we specified a model with [1,1] structure, representing the split of the two focal populations from a single ancestral population. Because our *Tangara* species have very high levels of overall population connectivity (see Results), we increased the maximum value of “M” (the proportion of migrants relative to the inferred ancestral population size) to 100 from the default value of 10. Additionally, we disabled dynamic population size changes by setting “Only sudden” equal to True. This reduces the number of parameters in the model by permitting only sudden changes in population sizes, as opposed to exponential or linear changes. We used a generation time of 1 year and a mutation rate of 2.04e-09 per site per generation, following precedent for other species in the tanager family (Lamichhaney et al. 2015). All other parameters were left at their default value. We ran 15 concurrent model searches for each pair of populations and extracted parameter values from the best-fitting model. Finally, we compared the net flux of migrants across each barrier in two ways: 1) the sum of the migration rates *m* (the proportion of migrants in a population each generation) for each population (i.e., *m_12_ + m_21_*), and 2) the sum of N\**m* for each population, representing the total number of individuals dispersing across a given barrier each generation (i.e., N_1_\**m_12_* + N_2_\**m_21_*). Because F_ST_ is tightly correlated with estimates of *m* (Figure 5) and alternative population groupings do not affect patterns of F_ST_ (Supplementary Figures 8-10), we only performed demographic modelling for the 23 population pairs delimited by the primary geographic population scenario described in Figure 1 and Table 2.

### Estimating genetic diversity and effective population size

We evaluated genetic diversity and effective population size at the species level using the subsampled autosomal dataset including all (variant and invariant) sites. First, for each sample, we calculated individual-level heterozygosity as one metric of species-level genetic diversity. We calculated individual heterozygosity from the 1D-SFS, which we created with ANGSD and winsfs as previously described. Next, we calculated species-level estimates of θ. Because θ=4N_e_μ at mutation-drift equilibrium, and our 8 study species are closely-related congeners that likely possess similar neutral mutation rates, we use θ as our proxy of effective population size, N_e_. Following the same general steps as before, we used ANGSD to generate the SAF across all samples within one species, and then created a 1D-SFS. We used the saf2theta and thetaStat functions to calculate two metrics of θ from the species-level 1D-SFS: 8_Watterson_, based on the number of segregating sites in the species, and 8_ν_, based on the pairwise nucleotide diversity. Both estimates of θ resulted in extremely similar patterns between biogeographic region (Supplementary Figure S12), so we report only 8_Watterson_ as our proxy of N_e_ here. Finally, we evaluated the relationship between 8_Watterson_ and range size using published geographic range sizes from BirdLife International.

## Results

### Bioinformatics

We performed 150 bp paired-end sequencing for N=166 individuals from our 8 study species across two lanes of a NovaSeq X. After adapter trimming and polyG removal, the average number of bases per individual decreased from 9.1 ± 2.3 billion to 7.9 ± 2.1 billion Q30 bases (range 1.6 – 12.4 billion bases). We aligned samples to the reference genome of *Camarhynchus parvulus* (GCA_901933205), another tanager, and removed overlapping and duplicate reads (average duplication rate 6.5% ± 2.4%). After excluding N=5 individuals with low mapping rates, we retained N=161 individuals for further analysis, which had an average mapping rate of 86.8% ± 6.3%. Finally, after realigning around indels, average per-base coverage across all samples was 5.6x ± 1.5x (range 1.7x – 8.6x). Species-level bioinformatic results are given in Table 1, and individual bioinformatic statistics are given in Supplementary Table S1.

### Intraspecific population structure

We assessed overall levels of intraspecific population structure within Andean and Amazonian species using principal components analyses (PCAs), the strength of isolation-by-distance (IBD), and admixture analyses. PCAs for Andean species were characterized by more distinct clusters corresponding to geographic populations, while PCAs for most Amazonian species showed U-shaped clines indicative of continuous variation across the landscape (Novembre and Stephens 2008) (Figure 2; Supplementary Figure S1). Furthermore, individuals accumulate genetic differences across space more quickly in the mountains than in the lowlands. The slope of the regression of IBD (β_IBD_) was significantly greater in Andean species than in Amazonian ones (Figure 3; Wilcoxon t-test p=0.029), and this is supported by higher average Mantel statistics (Andean average r=0.68; Amazonian average r=0.48) (Supplementary Table S3). Nevertheless, discrete structure is low overall, as expected for canopy frugivores like *Tangara.* Admixture analyses from PCAngsd supported an optimal number of K=2 ancestry groups for all 8 species (Supplementary Figures S2-S3). DAPC analyses also supported K=2 for most species except for two Andean taxa (*T. nigroviridis* and *T. parzudakii*), for which the lowest BIC was observed at K=3. Together, these results suggest that relatively subtle intraspecific population structure in these highly dispersive species is measurably greater in the Andes mountains than in the Amazonian lowlands.

**Figure 2.**
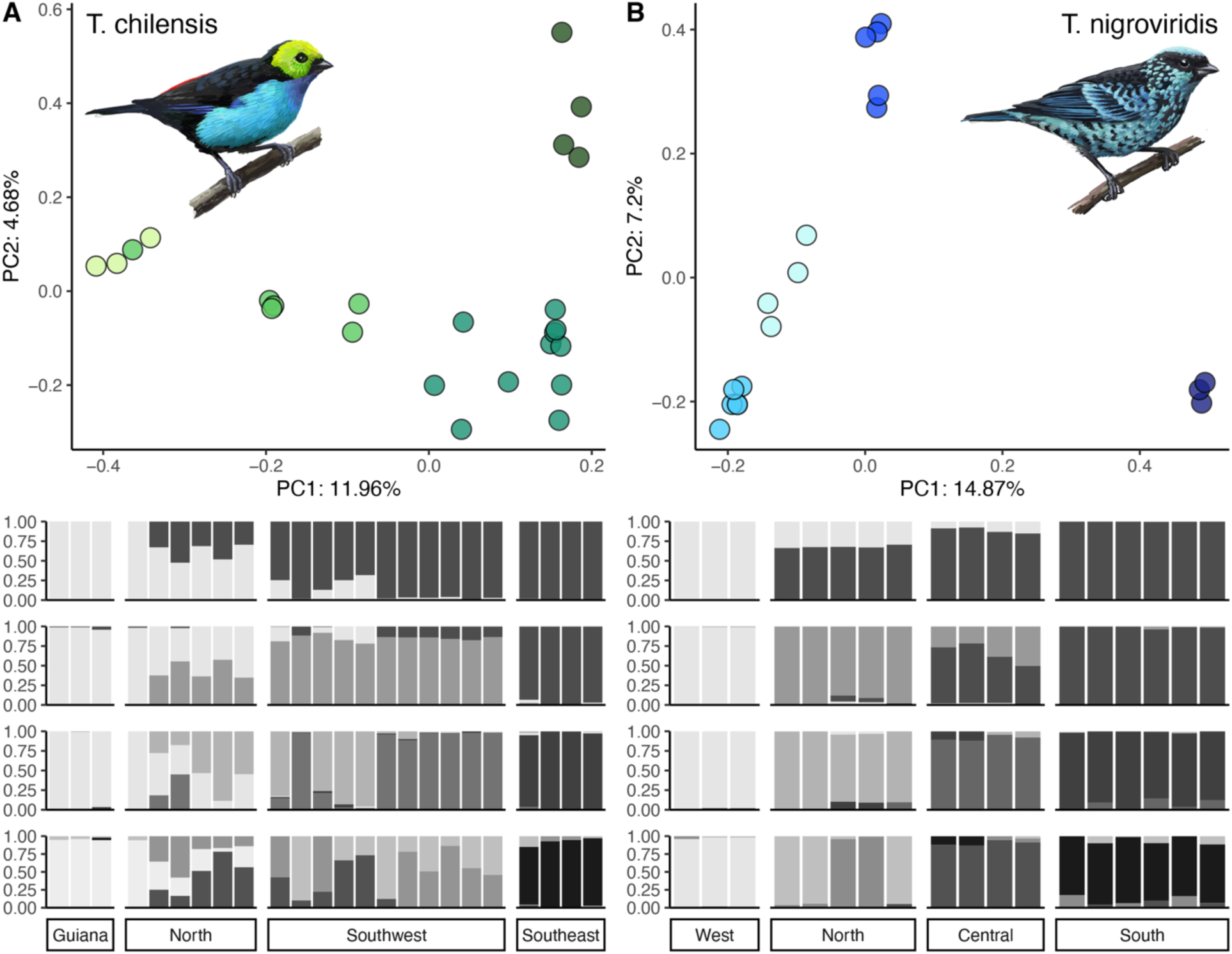
PCA and admixture results from PCAngsd for two example species: A) *T. chilensis*, an Amazonian species, and B) *T. nigroviridis*, an Andean species. Sample colors in the PCA correspond to the geographic populations described in Figure 1. Admixture barplots are for K=2-5, in ascending order. PCA and admixture analyses for all species are given in Supplementary Figures S1-S3. Illustrations by John Megahan.

**Figure 3.**
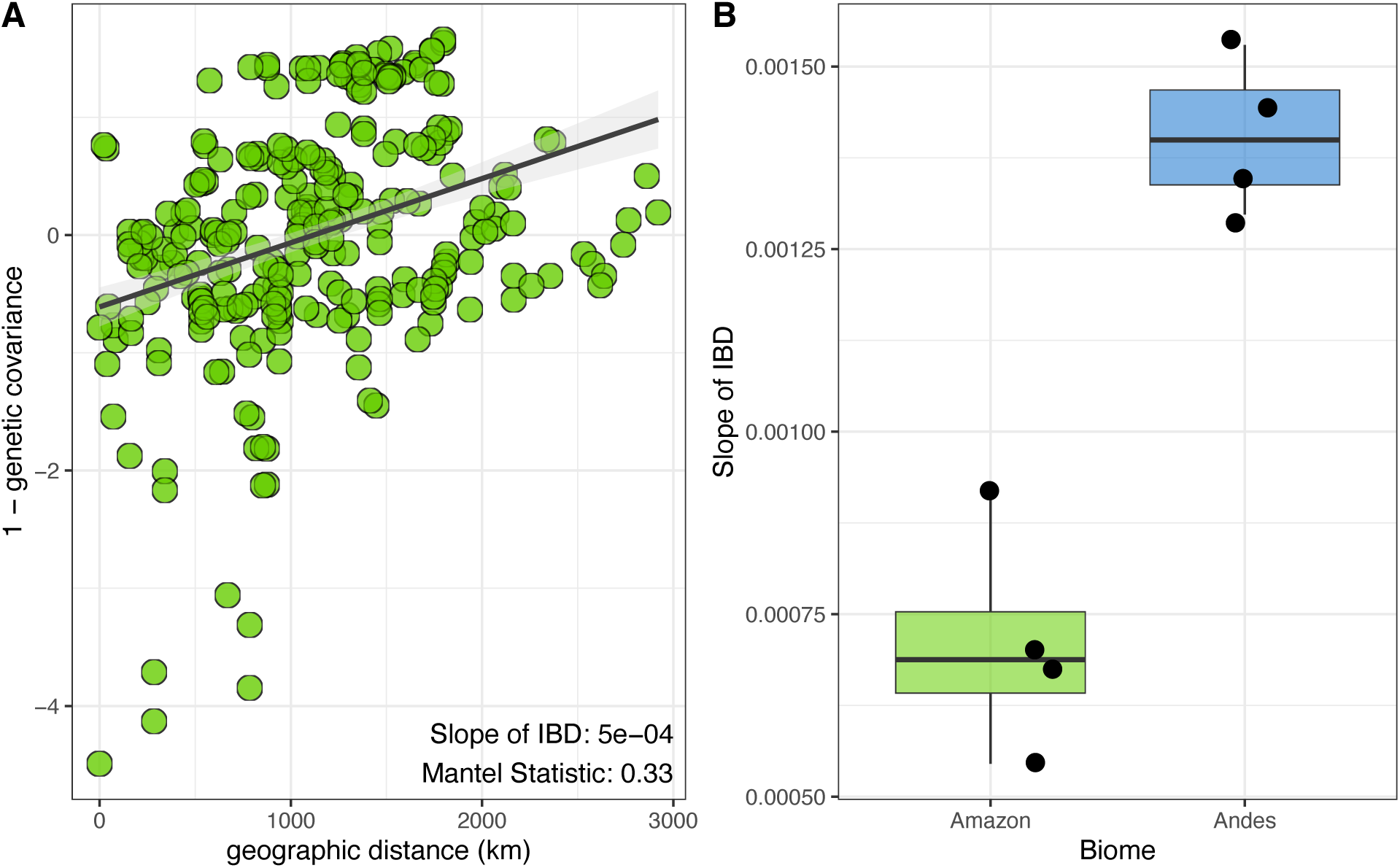
A) Example linear regression of the decay in pairwise genetic similarity against geographic distance for *T. chilensis*. Genetic similarity (1 – genetic covariance) was centered and z-transformed for all species. B) The strength of continuous population structure (β_IBD_) is significantly greater for Andean montane species than Amazonian lowland species. IBD regressions and Mantel statistics for all species are given in Supplementary Figures S4-S5.

Intraspecific population structuring revealed by the whole-genome data is not readily observed in the mitogenomes. Maximum likelihood phylogenetic trees from the full mitogenomes are characterized by extremely shallow divergences between tips, low nodal bootstrap support, and polyphyletic geographic populations (Supplementary Figures S6-S7). GMYC analyses on ultrametric ND2 gene trees did not reject panmixia for 7 of 8 species; that is, ND2 gene trees support just 1 interbreeding population for most *Tangara* species, compared to a median of 6 intraspecific populations across a diverse community of other Neotropical birds (Wacker and Winger 2024).

### Relative strength of geographic barriers

We evaluated the strength of individual geographic barriers in the Amazonian lowlands and Andean mountains by comparing levels of genetic differentiation and gene flow across them. Population pairs separated by a particular barrier in the Andes mountains had significantly higher values of genome-wide F_ST_ on average than population pairs in the Amazonian lowlands (Wilcoxon t-test p=0.0013; Figure 4). Indeed, Amazonian populations often had negative or 0 values of F_ST_, indicating homogeneity across a barrier (Figure 4; Supplementary Table SS3). One outlier pair of Andean populations—the West and Northeastern populations of *T. parzudakii*, corresponding to phenotypically-distinguished subspecies (O’Malley and Burns 2020)—exhibited markedly higher differentiation than other pairs (F_ST_=0.371). Nevertheless, average population differentiation across barriers remained significantly greater in the Andes even if this outlier is removed (p=0.0026). Furthermore, results were insensitive to alternative definitions of geographic populations (Supplementary Figures S8-S10). Naturally, there was some heterogeneity in the overall efficacy of different barriers and how they affected different species. For example, in the Andes, populations separated on different slopes in the Northern Andes and by the Huallaga river valley in Southern Peru tended to have higher values of F_ST_ than populations separated by other Andean barriers, while the lower stretch of the Amazon River tended to produce slightly higher F_ST_ values in the lowlands. However, we lack sufficient sampling to make general claims about the strengths of single barriers within each region.

**Figure 4.**
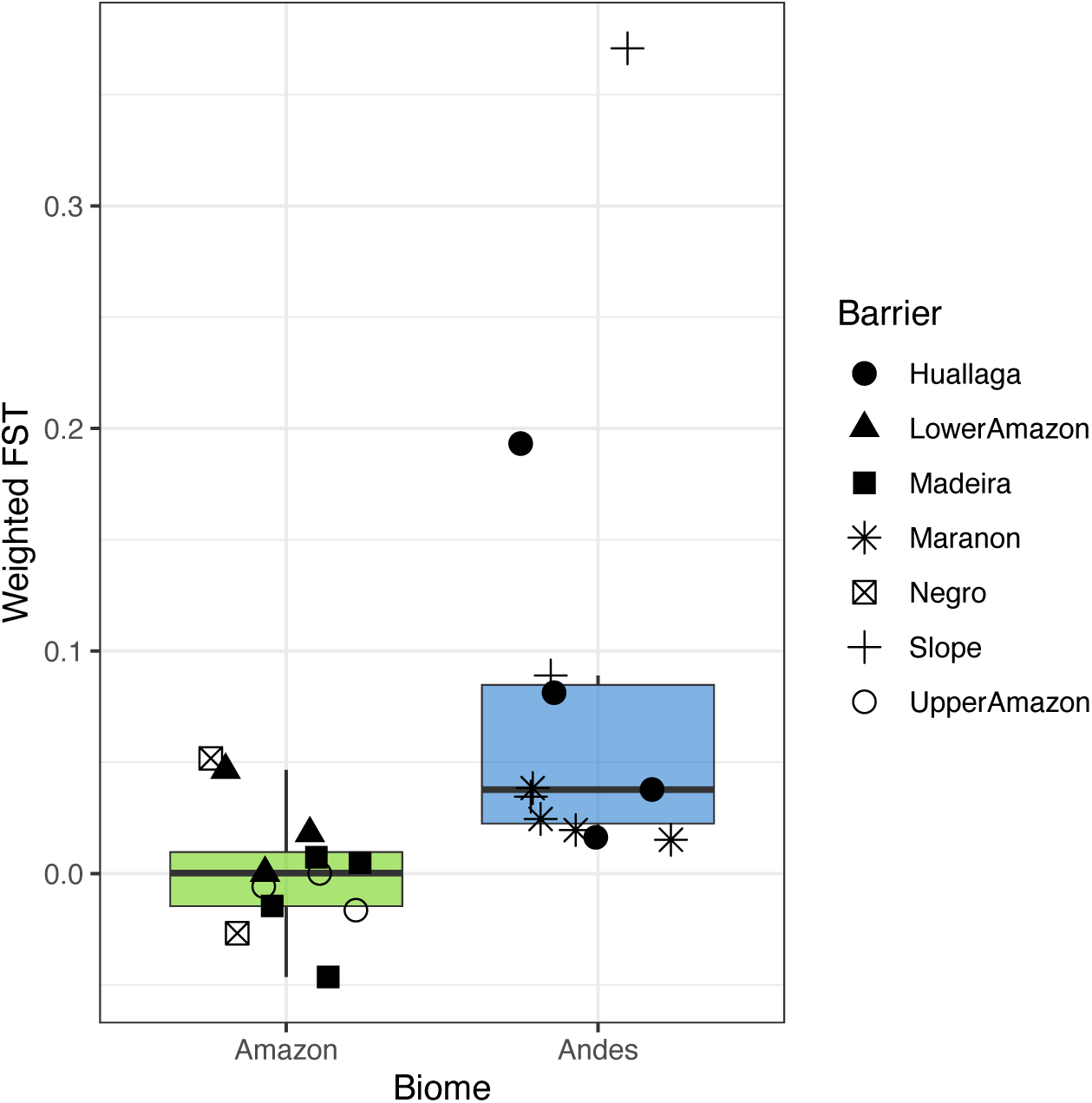
Pairwise population differentiation (F_ST_) across individual barriers in the Andes is significantly higher on average than in the Amazonian lowlands. The same statistically-supported difference is recovered for all alternative geographic population groupings (Supplementary Figure S10).

We also estimated levels of gene flow across barriers with demographic modeling of population pairs separated by individual barriers in GADMA (Noskova et al. 2023). The net flux of gene flow across barriers was higher on average in the Amazonian lowlands, when gene flow is measured both as the proportion of migrants in each population per generation (*m*; p=0.079) and as the total number of individuals dispersing across a barrier per generation (N*\*m;* p=0.00169), although only the latter metric reaches statistical significance (Supplementary Figure S11; Supplementary Table S4). The total number of individual migrants dispersing across a given barrier per generation remained significantly higher in the Amazonian lowlands even after removing 3 outliers with particularly high N\**m* values (p=0.0125; Figure 5A). Estimates of gene flow from GADMA showed a clear negative relationship with empirical F_ST_ across barriers: low values of F_ST_ are associated with very high numbers of migrants, and migration rates decay sharply to generate higher values of F_ST_ (Figure 5B).

**Figure 5.**
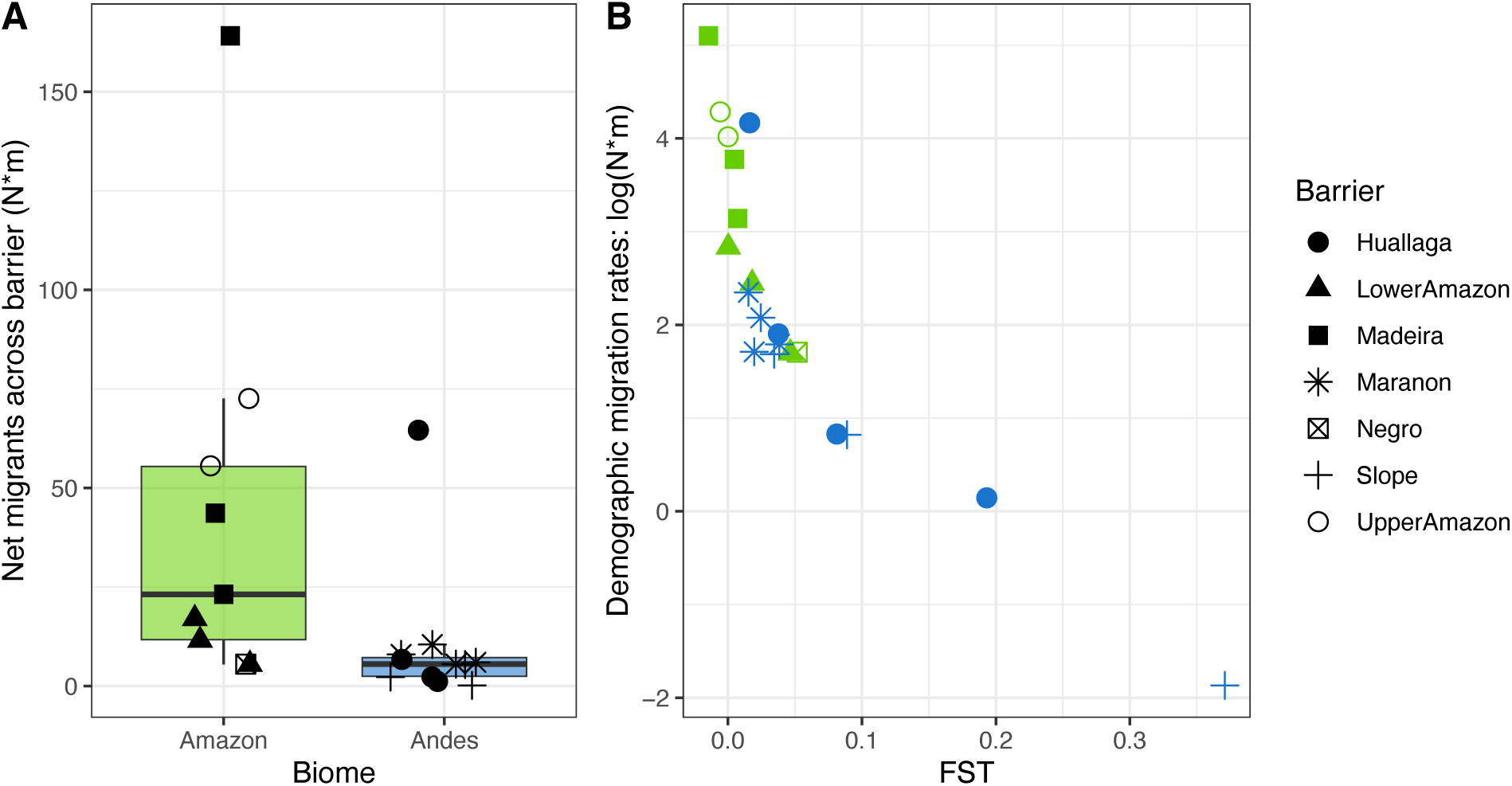
A) Net gene flow across particular barriers (presented as the total flux of individual migrants, N*m), inferred from the best fitting demographic model in GADMA. Three Amazonian outliers with exceptionally high parameter values are excluded from the comparison. Gene flow is significantly higher across Amazonian barriers than Andean barriers on average. B) The net rate of gene flow across a given barrier as estimated by demographic models decreases sharply as the empirically-fit value of F_ST_ across the same barrier increases. Number of migrants on the y-axis has been log-transformed for visual clarity.

### Genetic diversity and effective population size

We evaluated levels of genetic diversity for each species to test the hypothesis that smaller geographic ranges in the mountains lead to lower effective population sizes compared to the lowlands. Consistent with our predictions, every Andean species had significantly lower levels of individual heterozygosity than every Amazonian species (adjusted Wilcoxon’s p-values for all 16 species comparisons range from 3.1e-11 to 2.8e-6; Figure 6A). Species-level estimates of 8_Watterson_, our proxy of N_e_, were similarly much lower in the mountains than in the lowlands (p=0.0286; Figure 6B), as was 8_ν_ (Supplementary Table S4; Supplementary Figure S12). Furthermore, 8_Watterson_ appeared to positively correlate with geographic range size across these 8 species (Figure 6C), although this relationship was non-significant (R^2^=0.3875, p=0.099), likely due to small sample size. Nevertheless, this lends credence to the theoretical prediction that smaller geographic distributions characteristic of the Andes mountains lead to lower N_e_ and stronger genetic drift in montane taxa than Amazonian lowland taxa.

**Figure 6.**
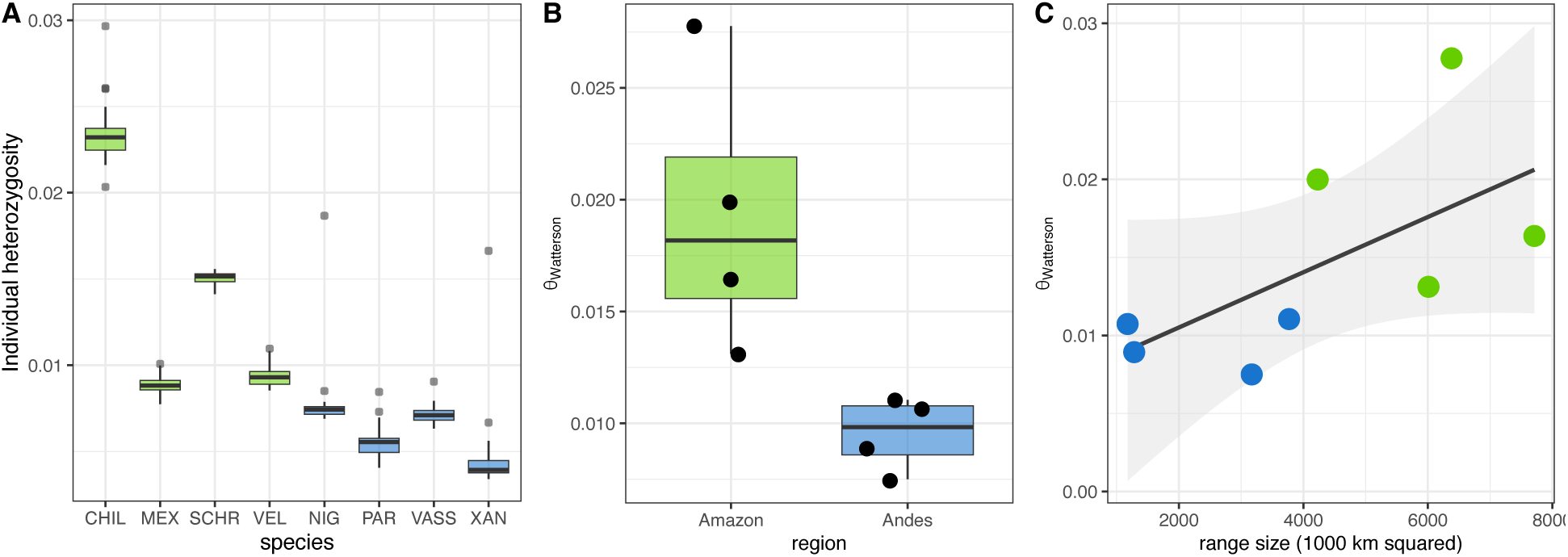
A) Boxplots of individual-level heterozygosity for each species. All Amazonian species have significantly greater individual-level heterozygosity than all Andean species. B) Species-level θ_Watterson_, a proxy for N_e_, is significantly greater in Amazonian species than Andean species. C) Data for our eight study species suggests a positive relationship between θ_Watterson_ and range size, wherein the four species with ranges above 4000 km^2^ have higher genetic diversity than the four species with smaller range size, but the linear relationship is non-significant.

## Discussion

In this study, we evaluated the phylogeographic divergence dynamics of Andean mountain versus Amazonian lowland birds. Using 161 moderate-coverage whole genome sequences across 8 species, we showed that ecologically-similar *Tangara* tanagers have relatively greater signals of population structuring in the mountains than in the lowlands. Nevertheless, levels of discrete structure were low overall, consistent with expectations for extremely dispersive canopy frugivores in humid tropical forests. Across individual geographic barriers, Andean species showed higher levels of population differentiation and lower levels of gene flow than Amazonian species, supporting our hypothesis of stronger barrier-driven allopatry in the mountains compared to the river-structured lowlands. Additionally, Andean species were characterized by lower levels of genetic diversity and smaller effective population sizes than Amazonian species, likely due to their smaller geographic distributions. These results together suggest that the initial stages of speciation occur more readily in the Andes than in the Amazon, owing to the combined effects of stronger physical barriers and smaller effective population sizes in the mountains.

### Population structuring in the mountains versus lowlands

Andean *Tangara* tanagers show elevated levels of population structuring relative to their Amazonian counterparts. Principal component and admixture analyses reveal tighter geographic clustering of individuals from the mountains than the lowlands (Figure 2; Supplementary Figures S1-S3), and continuous genetic differentiation (isolation-by-distance) accumulates more rapidly in the mountains (Figure 3; Supplementary Figures S4-S5). However, the amount of discrete population structure in the canopy frugivore *Tangara* is quite low overall, with admixture analyses and K-means clustering supporting between 2 (most species) and 3 (two Andean species as evaluated with DAPC) ancestry groups. Additionally, whole mitogenomes display extremely shallow phylogenetic structure and high polyphyly with respect to geographic populations (Supplementary Figures S6-S7). Thus, our results support the general elevational gradient in phylogeographic diversity in Neotropical birds (Wacker and Winger 2024), but this is evident only with more powerful analyses utilizing our whole-genome datasets. This illustrates the power and importance of genomics for uncovering evidence of subtle population genetic divergence (McGaughran et al. 2022, Kimmitt et al. 2024), especially for highly-dispersive taxa like *Tangara*.

### Barrier efficacy in the mountains versus the lowlands

We hypothesized that greater phylogeographic structure in the mountains could be driven by relatively more effective physical barriers. Supporting this hypothesis, we found that individual geographic barriers in the Andes promoted greater population differentiation (F_ST_) than those in the Amazonian lowlands, where species often showed genomic homogeneity across barriers (Figure 4). Additionally, the number of individuals that cross a barrier each generation (N*m), as estimated with demographic modeling, was significantly lower on average in the mountains (Figure 5A). For the same geographic barrier and pair of populations within a species, the inferred rate of migration decreases sharply as F_ST_ increases, mirroring classic population genetic relationships between N*m and population divergence (Figure 5B; Wright 1931). Simulation studies show that the effects of physical barriers on population divergence become more confounded by additional factors like historical demography and selection when taxa have low dispersal capabilities (Araya-Donoso et al. 2022); thus, our ability to detect these differential effects of regional barriers on genetic differentiation is likely enhanced by our study system.

Although we see a strong pattern in the average barrier efficacy across biogeographic regions, there is some heterogeneity in the level of divergence across different barriers and species within a single biogeographic region (Figure 4). For example, in Amazonia, populations of *Tangara* show little to no divergence across the Madeira River or across the upper course of the Amazon River in western Amazonia, but some species have structure across the Negro and lower course of the Amazon River. In the Andes, differentiation across the Marañón Valley tends to be relatively lower than divergence across the Huallaga Valley and across slopes of the mountain range, but the amount of divergence across these more effective barriers varies across species. Nevertheless, detailed comparisons of individual barriers and species’ responses within each biogeographic region (Andes or Amazonia) would require more samples with greater spatial resolution than currently available.

### Genetic diversity and effective population size

We found consistent and robust differences in genetic diversity between montane and lowland species. All Andean birds had significantly lower amounts of individual heterozygosity and smaller values of θ_Watterson_ (our proxy for N_e_) than all Amazonian birds (Figure 6A-B). Because the terrestrial surface area of the tropical Andes mountain range is considerably smaller than that of the Amazonian lowlands, neutral theory would predict lower genetic diversity and N_e_ in the mountains via the effect of range size on individual abundance (Lawton 1993, Gaston et al. 1997). Indeed, continuous range size appeared positively correlated with θ_Watterson_, although the linear relationship across our 8 study species is marginally non-significant (Figure 6C). Because species with lower levels of genetic diversity should experience more rapid population differentiation across a barrier (Charlesworth 2009, Bromham 2020), it is likely that these intrinsic genetic differences between Andean and Amazonian species are contributing to the elevated values of F_ST_ across barriers characterizing montane taxa.

Although there are strong reasons to expect lower genetic diversity in species with smaller geographic range sizes (Leffler et al. 2012, Ellegren and Galtier 2016), empirical evidence has been mixed. For example, Romiguier et al. (2014) reported a strong influence of life history strategy but no role for geographic range on patterns of species-level genetic diversity. However, this study included widely divergent species from across the animal tree of life, and reported an extremely strong taxonomic effect at the family level; similarly, a high degree of phylogenetic signal in genetic diversity across metazoans was found by Buffalo (2021). Here, by restricting our analysis to closely-related congeners with similar life history, body size, and ecological traits, we were able to recover a role for geographic distribution on genetic diversity. A similar relationship has been corroborated in other bird systems, with evidence for a positive impact of range size in island versus continental species (Leroy et al. 2021) and of geographic extent of historical suitable habitat in North American species (Brüniche-Olsen et al. 2021) on genetic diversity and N_e_. Nevertheless, the predicted effect of range size on genetic diversity is through its relationship with census population size, which itself is mediated by population density. Thus, variation in species’ abundance could attenuate the predicted correlation between range size and diversity.

### Mechanisms underlying Andean-Amazonian biodiversity gradients

Like global latitudinal gradients in different metrics of biodiversity (e.g., Rabosky et al. 2018, Harvey et al. 2020), there seems to be multiple contrasting elevational gradients in biodiversity patterns characterizing the megadiverse avifauna of the Andes-Amazonia system. First, species richness in Neotropical birds is highest in the Amazonian lowlands, and decreases with elevation up the slope of the adjacent Andes mountain range (Terborgh et al. 1984, Kattan and Franco 2004, Quintero and Jetz 2018). However, there is some evidence that contemporary rates of species formation in birds are reduced in the Amazonian lowlands and highest in the Andes mountains (Weir 2006, Quintero and Jetz 2018). Furthermore, levels of mitochondrial phylogeographic diversity also follow an elevational gradient, with Andean birds having more intraspecific population structure than Amazonian birds (Wacker and Winger 2024). Here, we uncover different dynamics of phylogeographic divergence that can potentially reconcile these conflicting trends in species richness on the one hand and the formation of incipient species or full species on the other.

Our data suggest that small population sizes and less permeable geographic barriers drive stronger allopatry and population differentiation in Andean birds compared to Amazonian ones. This enhanced propensity for populations to fragment and diverge from one another ought to initiate the process of speciation (Tobias et al. 2020) more readily in the mountains than in the lowlands, resulting in more phylogeographic diversity and higher tip-level speciation rates. However, these types of metapopulations (smaller and more fragmented) are also more prone to demographic extinction (Levins 1969), raising the possibility that young populations and species aren’t persisting through evolutionary timescales in a way that generates long-term species diversity. Indeed, it has been previously suggested that many incipient or young species are ephemeral across evolutionary timescales (Rosenblum et al. 2012, Harvey et al. 2017b, Singhal et al. 2018, Wacker and Winger 2024), and failure to persist can be a major control on the build-up of biodiversity (Harvey et al. 2019). In contrast, populations in the Amazonian lowlands tend to be larger and more connected. While this connectivity may dampen the formation of incipient species *relative to mountain taxa*, such metapopulations tend to be more stable and persistent through time, which could better foster the long-term persistence of species-rich assemblages.

### Limitations and future directions

We conducted this study using 8 closely-related and ecologically comparable tanagers. While this allowed us to isolate the effects of different biogeographic regions on phylogeographic dynamics, it remains unclear whether our recovered patterns of structure, population differentiation, and genetic diversity would be borne out in other Neotropical taxa. Despite the closely-interconnected geoclimatic histories (Hoorn et al. 2010, Bicudo et al. 2019) and species assemblages (Brumfield and Edwards 2007, van Els et al. 2021, Pérez-Escobar et al. 2022) of the Andes mountain system and the adjacent Amazonian lowlands, very few studies have assessed how microevolutionary processes play out within each region using a common methodological framework. We encourage future comparative phylogeographic work bridging the Andes-Amazonia system in order to evaluate the generality of the diversification scenario presented here.

In this work, our comparative sampling regime restricts us to a coarse analysis of genomic divergence across enormous geographic areas. Trade-offs among sequencing breadth across the genome, depth of coverage, and number of samples are inherent to evolutionary genomics (Lou et al. 2021), and this problem is compounded when comparing evolutionary trajectories across multiple species and/or regions. More thorough sampling regimes provide enhanced resolution of the nuanced effects of complex Neotropical landscapes on population divergence histories (e.g., Berv et al. 2021, Musher et al. 2022, Moncrieff et al. 2024), but these are often restricted to the best-sampled species. Indeed, sparse sampling of the immense diversity contained in hard-to-access Neotropical regions is a major limitation to our understanding of biotic diversification in the Andes-Amazonia region (Hortal et al. 2015, Ribas et al. 2022), especially given the known impact of spatial sampling regimes on population genetic inference (Battey et al. 2020). Together, this underscores the importance of growing and maintaining accessible natural history collections (Nachman et al. 2023) and open biodata repositories (Leigh et al. 2021) for more refined study of ecological and evolutionary processes in natural systems.

### Conservation implications

Organisms can respond to global climate change by in situ adaptation to altered environmental conditions or by shifting their ranges to track their niches, and otherwise face local extirpation or range-wide extinction (Davis et al. 2005). Tropical mountain birds have demonstrated rapid upslope range shifts as mean annual temperatures increase (Freeman and Freeman 2014), including in the tropical Andes mountains (Freeman et al. 2018). This puts tropical mountain communities on an “escalator to extinction,” whereby species near the summits of their mountain peaks may disappear because they lack additional upslope habitat to shift into (Sekercioglu et al. 2008). In theory, this extirpation scenario could be averted if species were able to adapt in place to increased temperatures and modified biotic interactions (for examples of climate change adaptation, see Bi et al. 2019, Turbek et al. 2023). Genome-wide genetic variation is crucial to the adaptive capacity of organisms, and is particularly important in a conservation context (Lande and Shannon 1996, Kardos et al. 2021, Mathur et al. 2023). Here, our results suggest strong disparities in the standing levels of genetic diversity characterizing tropical mountain versus lowland taxa, with Andean species having considerably less genome-wide variation and thus presumably reduced capacity to adapt to rapid climate change. This emphasizes the importance and urgency of curbing anthropogenic climate change, especially for the conservation of tropical mountain species (Parmesan 2006).

## Acknowledgements

For providing tissue samples, we thank the curators, collections staff, and field collectors from the following institutions: Louisiana State University Museum of Natural Sciences, Academy of Natural Sciences of Drexel University, American Museum of Natural History, Field Museum of Natural History, Harvard University Museum of Comparative Zoology, University of California Berkeley Museum of Vertebrate Zoology, University of Kansas Biodiversity Institute and Natural History Museum, University of New Mexico Museum of Southwestern Biology, and Smithsonian Museum of Natural History. We thank Teresa Pegan and Abby Kimmitt for invaluable bioinformatics training. We also thank Nicole Adams, Zach Hancock, Gideon Bradburd, and Robb Brumfield for discussion and feedback. Next-generation sequencing for this project was carried out in the Advanced Genomics Core at the University of Michigan, and computational resources and support were provided by the Advanced Research Computing group at the University of Michigan’s Technology Services.

