## Supplementary Tables S2-S5; Supplementary Figures S1-S12 for "Distinct intraspecific diversification dynamics in Neotropical montane versus lowland birds revealed by whole-genome comparative phylogeography"

**Supplementary Table S2.** Mitogenome assembly with NOVOplasty and results. A conspecific ND2 sequence from GenBank was used as the seed for mitogenome assembly runs in NOVOplasty for each sample. The average and standard deviation of mitogenome sequencing coverage for each species is given.

| **Species** | **ND2 seed (Accession)** | **ND2 seed (Specimen)** | **Average coverage** |
| --- | --- | --- | --- |
| T. chilensis | EU648058 | MVZ FC21600 | 6032 ± 4654 |
| T. mexicana | EU648079 | LSUMNS B18465 | 8215 ± 6640 |
| T. nigroviridis | EU648082 | LSUMNS B1627 | 8110 ± 5487 |
| T. parzudakii | EU648084 | LSUMNS B30007 | 7827 ± 6100 |
| T. schrankii | EU648090 | LSUMNS B34932 | 7288 ± 4830 |
| T. vassorii | EU648093 | LSUMNS B1711 | 6141 ± 4960 |
| T. velia | EU648094 | FMNH 390060 | 9840 ± 7987 |
| T. xanthocephala | EU648097 | LSUMNS B34922 | 8550 ± 6093 |

**Supplementary Table S3.** Isolation-by-distance analyses for each species. The strength of IBD is captured with the slope of the linear regression of pairwise 1 – genetic covariance versus geographic distance (“lm_slope”). We also report the mantel statistic giving the correlation between the pairwise genetic covariance matrix and the geographic distance matrix (“mantel_r). Significance key: * p<0.05; ** p<0.01; ***p<0.001

| **Species** | **lm_slope** | **mantel_r** |
| --- | --- | --- |
| T. chilensis | 0.00054518 *** | 0.33285649 *** |
| T. mexicana | 0.00090957 *** | 0.67316317 *** |
| T. nigroviridis | 0.0014484 *** | 0.63534117 *** |
| T. parzudakii | 0.00152946 *** | 0.64034635 *** |
| T. schrankii | 0.0007023 *** | 0.37640121 *** |
| T. vassorii | 0.0013503 *** | 0.71600658 *** |
| T. velia | 0.00067299 *** | 0.55266525 *** |
| T. xanthocephala | 0.00129722 *** | 0.73205194 *** |

**Supplementary Table S4.** FST and GADMA migration parameter estimates for each pair of geographic populations separated by a given barrier. Numbers in parentheses after each population name are the sample size for that population and species. FST is the weighted genome-wide (autosomal) divergence between populations calculated with the Bahtia estimator (-which 1) in ANGSD. Migration parameters are net flux across rivers; *m* is the sum of the unidirectional per-generation migration rates between populations (i.e., *m_12_ + m_21_*) and N**m* is the number of individuals crossing a barrier each generation (i.e., N_1_* *m_12_* + N_2_* *m_21_*).

| **Species** | **Biome** | **Pop1** | **Pop2** | **Barrier** | **FST** | **m**  **(rate)** | **N*m**  **(inds)** |
| --- | --- | --- | --- | --- | --- | --- | --- |
| T. chilensis | Amazon | Guiana (3) | North Amazon (6) | Negro | -0.02678 | 4.63e-05 | 2519.512 |
| T. chilensis | Amazon | Guiana (3) | Southeast Amazon (4) | Lower Amazon | 0.00026 | 2.90e-06 | 17.009 |
| T. chilensis | Amazon | North Amazon (6) | Southwest Amazon (11) | Upper Amazon | -0.00578 | 3.79e-05 | 72.583 |
| T. chilensis | Amazon | Southeast Amazon (4) | Southwest Amazon (11) | Madeira | -0.01454 | 3.99e-05 | 164.128 |
| T. mexicana | Amazon | Guiana (5) | North Amazon (3) | Negro | 0.0517 | 2.01e-06 | 5.510 |
| T. mexicana | Amazon | Guiana (5) | Southeast Amazon (6) | Lower Amazon | 0.04654 | 3.41e-06 | 5.465 |
| T. mexicana | Amazon | North Amazon (3) | Southwest Amazon (7) | Upper Amazon | 0.00007 | 5.72e-05 | 55.573 |
| T. mexicana | Amazon | Southeast Amazon (6) | Southwest Amazon (7) | Madeira | 0.0073 | 1.65e-05 | 23.149 |
| T. schrankii | Amazon | North Amazon (5) | Southwest Amazon (11) | Upper Amazon | -0.0164 | 6.82e-05 | 2543.408 |
| T. schrankii | Amazon | Southeast Amazon (2) | Southwest Amazon (11) | Madeira | -0.04649 | 7.22e-05 | 2525.676 |
| T. velia | Amazon | Guiana (6) | Southeast Amazon (3) | Lower Amazon | 0.01808 | 9.14e-06 | 11.540 |
| T. velia | Amazon | Southeast Amazon (3) | Southwest Amazon (10) | Madeira | 0.00488 | 2.22e-05 | 43.638 |
| T. nigroviridis | Andes | Central Andes (4) | Northeast Andes (5) | Maranon | 0.02455 | 1.16e-05 | 7.977 |
| T. nigroviridis | Andes | Central Andes (4) | South Andes (6) | Huallaga | 0.01621 | 1.43e-05 | 64.562 |
| T. nigroviridis | Andes | Northeast Andes (5) | West Andes (3) | Slope | 0.08892 | 3.79e-06 | 2.273 |
| T. parzudakii | Andes | Central Andes (4) | Northeast Andes (5) | Maranon | 0.0384 | 1.43e-05 | 5.996 |
| T. parzudakii | Andes | Central Andes (4) | South Andes (5) | Huallaga | 0.03768 | 9.24e-06 | 6.714 |
| T. parzudakii | Andes | Northeast Andes (5) | West Andes (5) | Slope | 0.37088 | 3.69e-07 | 0.155 |
| T. vassorii | Andes | Central Andes (2) | Northeast Andes (6) | Maranon | 0.01523 | 1.62e-05 | 10.471 |
| T. vassorii | Andes | Central Andes (2) | South Andes (10) | Huallaga | 0.08129 | 3.71e-06 | 2.292 |
| T. vassorii | Andes | Northeast Andes (6) | West Andes (2) | Slope | 0.03449 | 1.09e-05 | 5.398 |
| T. xanthocephala | Andes | Central Andes (4) | Northeast Andes (8) | Maranon | 0.01958 | 3.98e-05 | 5.541 |
| T. xanthocephala | Andes | Central Andes (4) | South Andes (8) | Huallaga | 0.19318 | 1.50e-06 | 1.156 |

**Supplementary Table S5.** Genetic diversity statistics for each species, and the breeding/resident range size from BirdLife International.

| **Species** | **θ**_π_ | **θ_Watterson_** | **Tajima’s D** | **Individual heterozygosity** | **Range size (km^2^)** |
| --- | --- | --- | --- | --- | --- |
| T. chilensis | 0.01973 | 0.02776 | -1.06835 | 0.02347 ± 0.00183 | 6380000 |
| T. mexicana | 0.01007 | 0.01637 | -1.45144 | 0.00892 ± 0.00060 | 7710000 |
| T. nigroviridis | 0.00874 | 0.01105 | -0.80896 | 0.00804 ± 0.00268 | 3770000 |
| T. parzudakii | 0.00739 | 0.00893 | -0.66802 | 0.00561 ± 0.00110 | 1282100 |
| T. schrankii | 0.01341 | 0.01999 | -1.25204 | 0.01506 ± 0.00040 | 4230000 |
| T. vassorii | 0.00845 | 0.01073 | -0.81114 | 0.00720 ± 0.00059 | 1176000 |
| T. velia | 0.01046 | 0.01312 | -0.77688 | 0.00939 ± 0.00066 | 6010000 |
| T. xanthocephala | 0.00538 | 0.00750 | -1.08252 | 0.00484 ± 0.00290 | 3170000 |

**Supplementary Figure S1.** Principal component analysis from ANGSD for all Amazonian (A-D) and all Andean (E-F) species. Geographic populations of samples are colored according to the legend in A and E, and match the population designations of Figure 1 in the main manuscript.

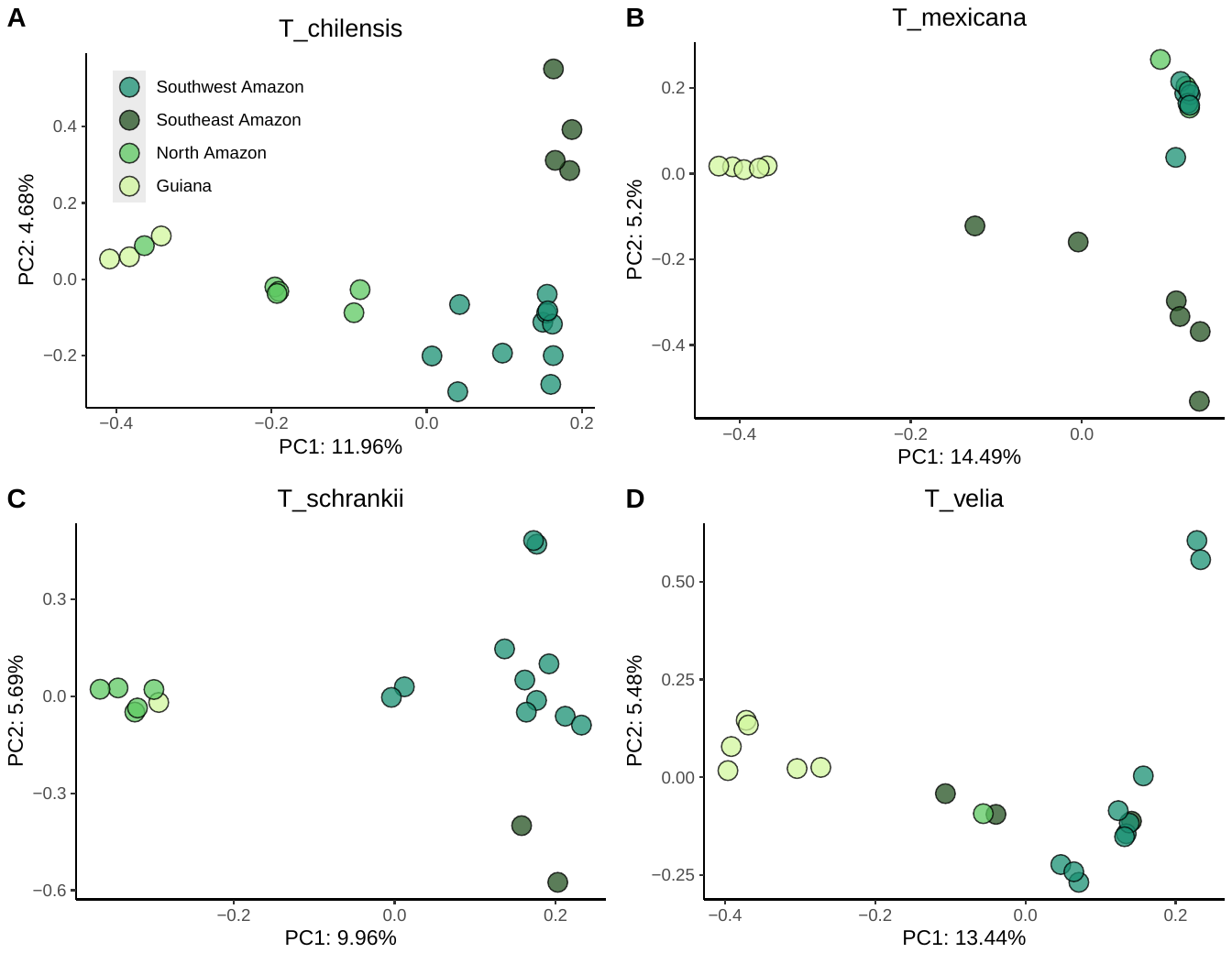

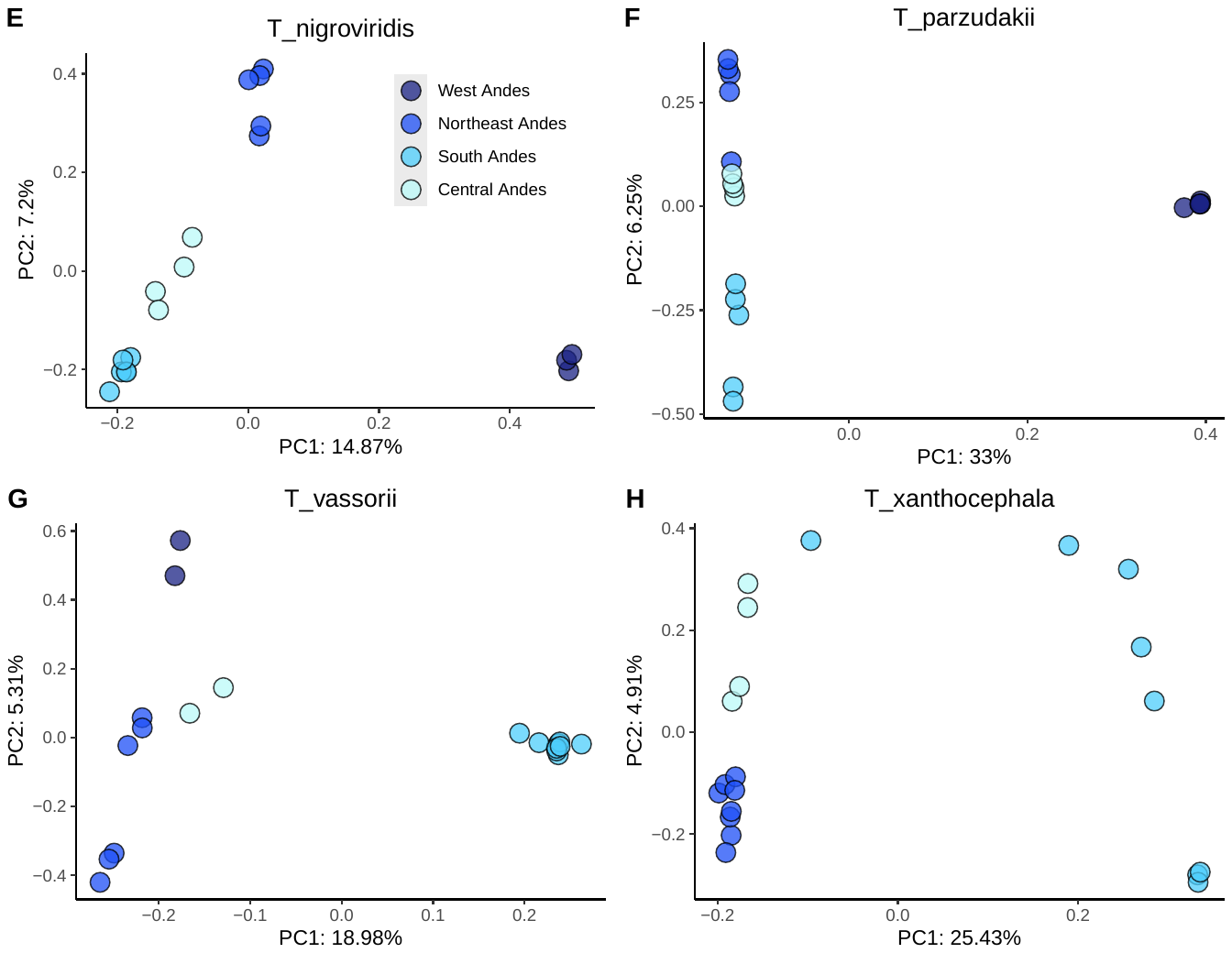

**Supplementary Figure S2.** Admixture analyses from ANGSD for all Amazonian species. The best-fitting K is chosen by default in pcangsd; K=2 was the best K for all species. Manual runs of K=3-5 are also shown for comparison.

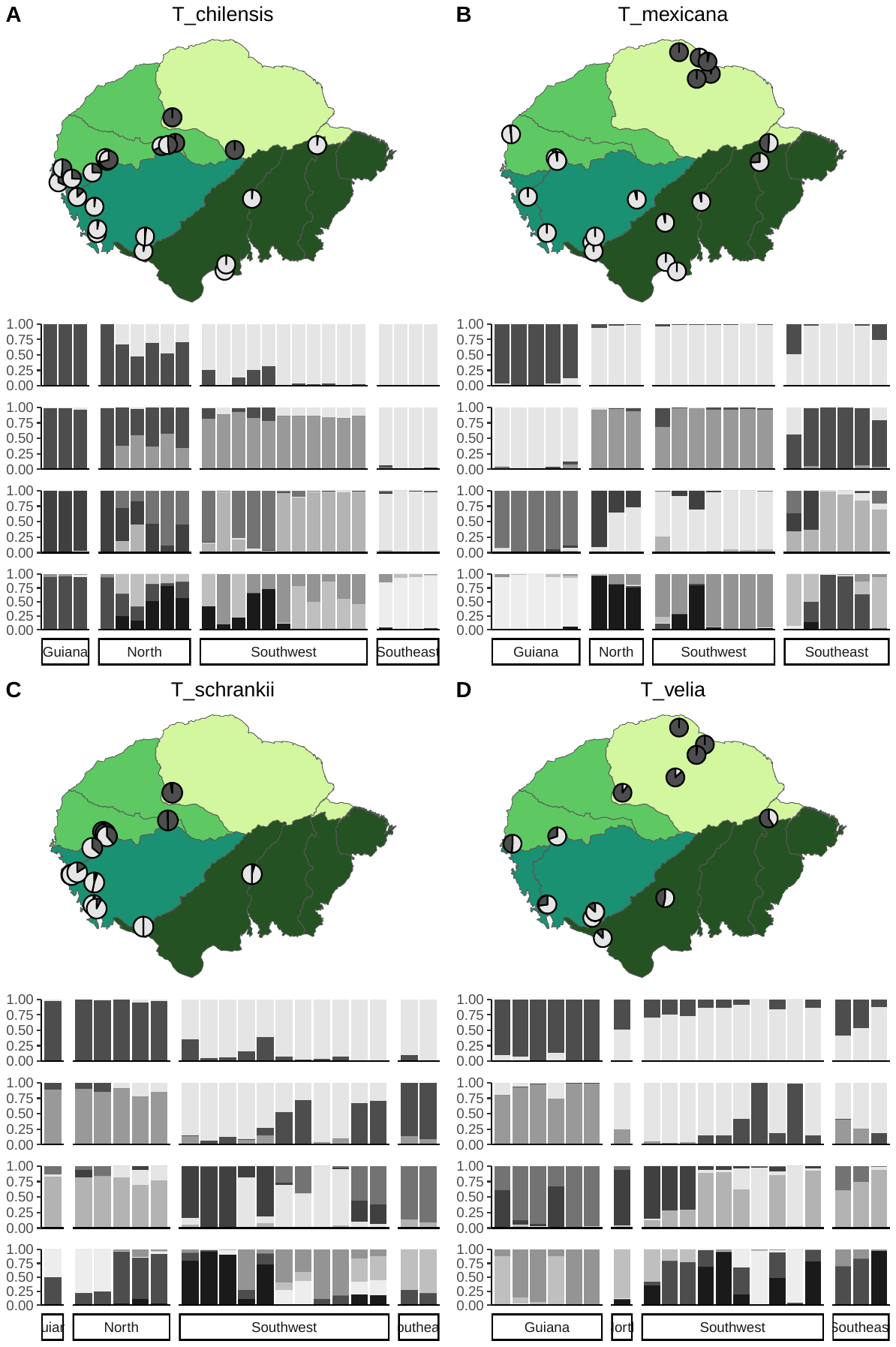

**Supplementary Figure S3.** Admixture analyses from ANGSD for all Andean species. The best-fitting K is chosen by default in pcangsd; K=2 was the best K for all species. Manual runs of K=3-5 are also shown for comparison.

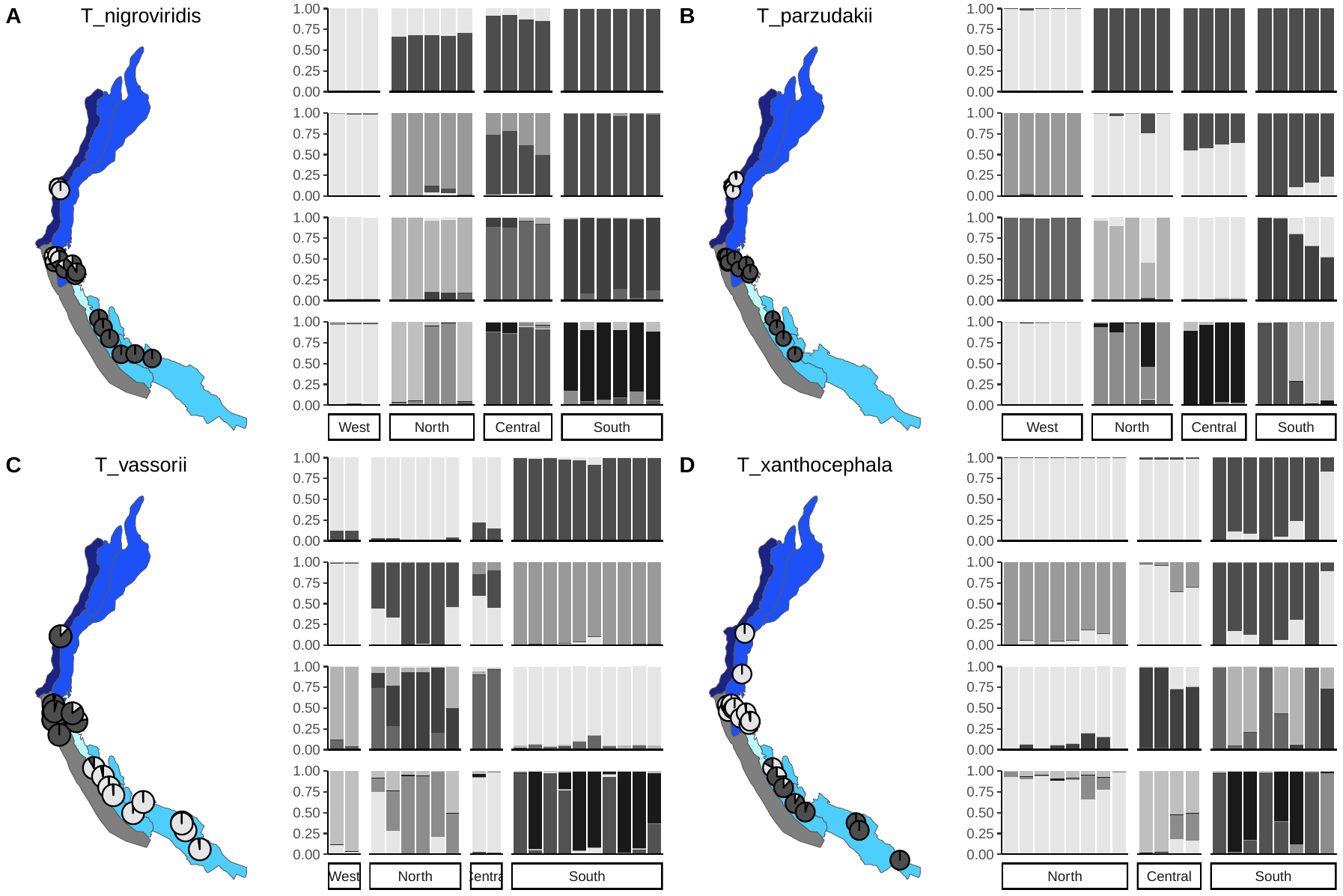

**Supplementary Figure S4.** Isolation-by-distance regression plots for each Amazonian species. The slope of the regression and the mantel statistic are reported in the bottom right corner.

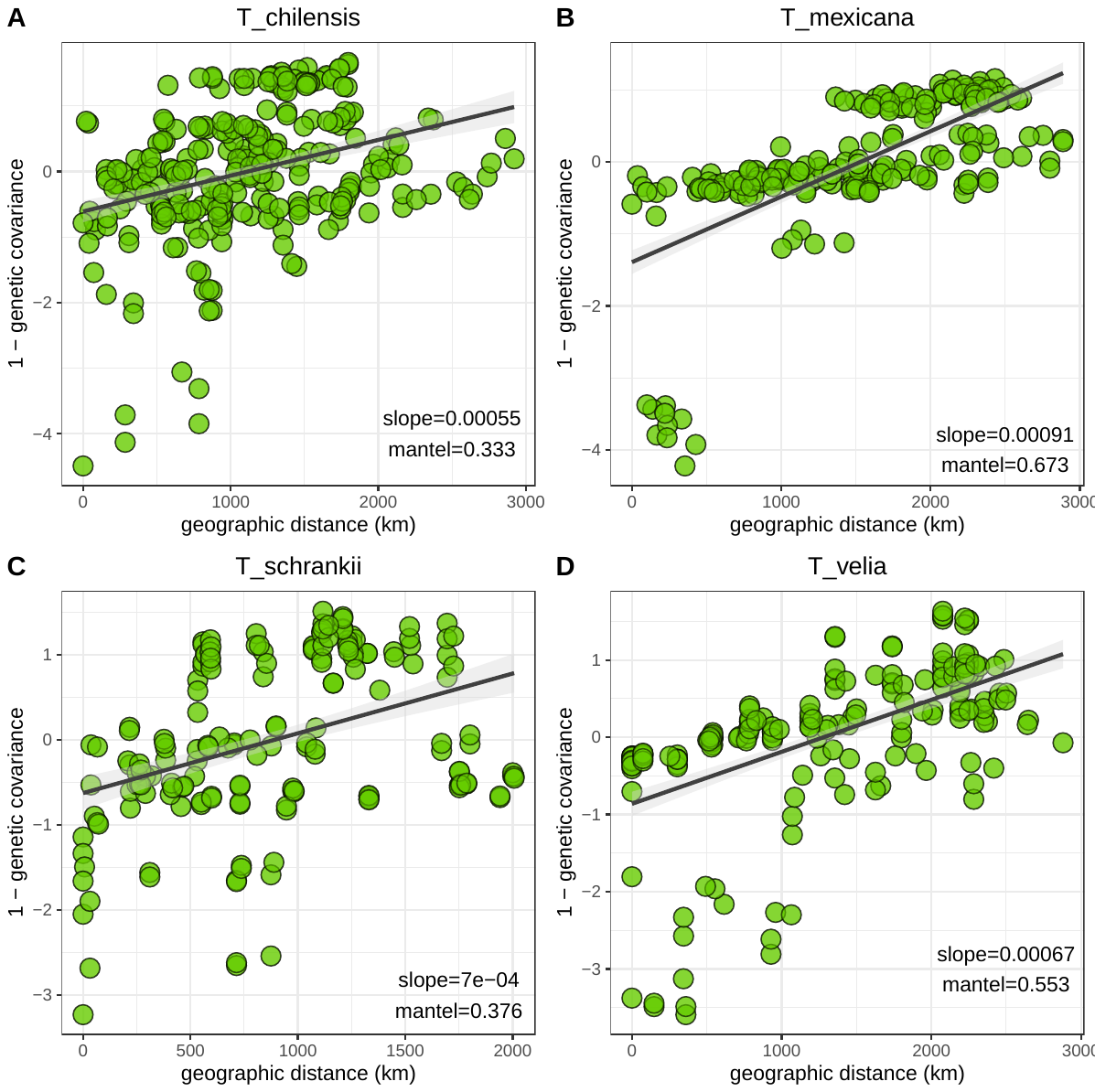

**Supplementary Figure S5.** Isolation-by-distance regression plots for each Andean species. The slope of the regression and the mantel statistic are reported in the bottom right corner.

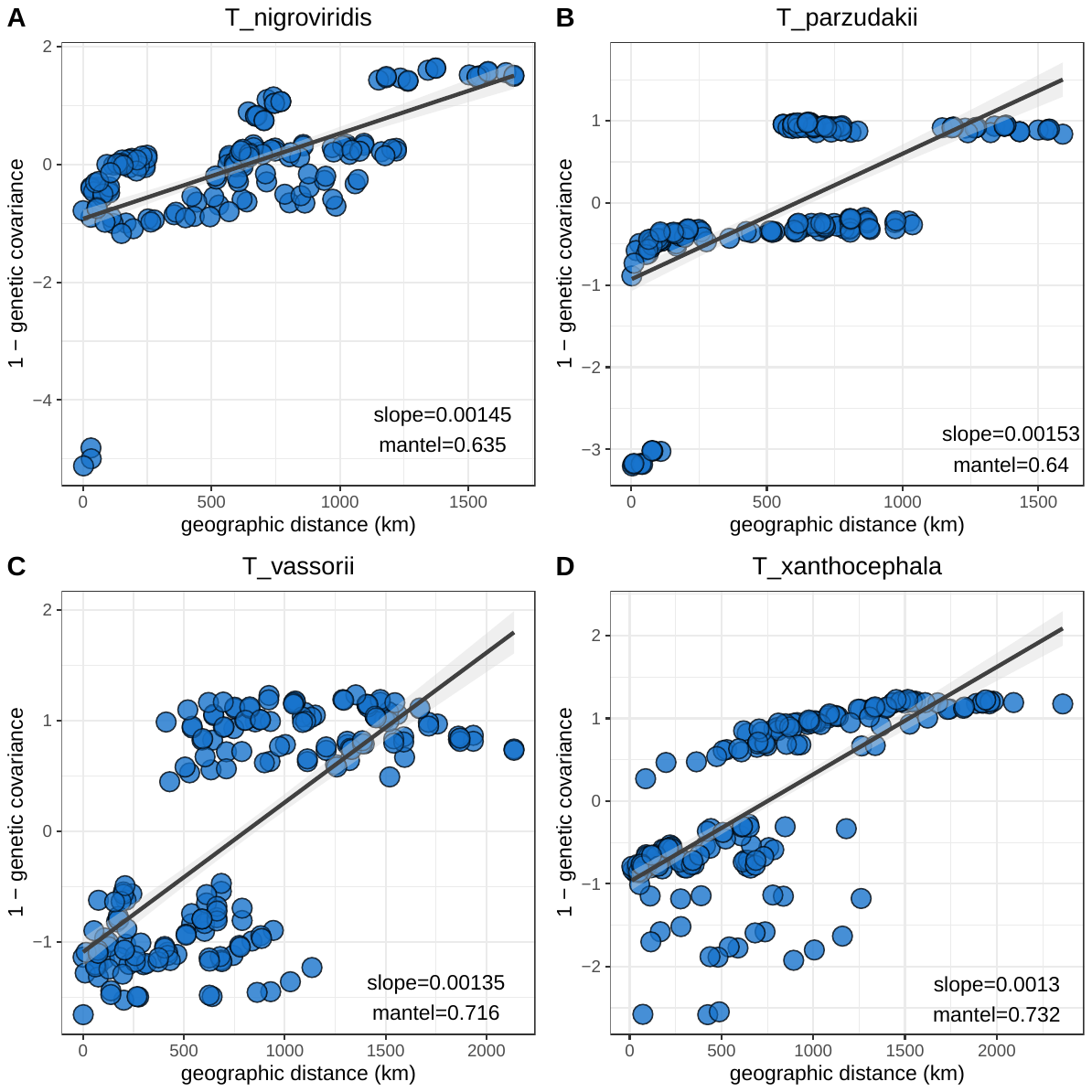

**Supplementary Figure S6.** Mitochondrial phylogenetic trees for the full 13 protein-coding gene sets for all Amazonian species. Trees were made with IQ-Tree 2 using automatic model selection from alignments partitioned by gene and codon, with 1000 ultrafast bootstrap replicates. Tips are color labeled according to their geographic population, as in Supplementary Figure 1. Nodes with > 80% bootstrap support are labeled with a solid black square.

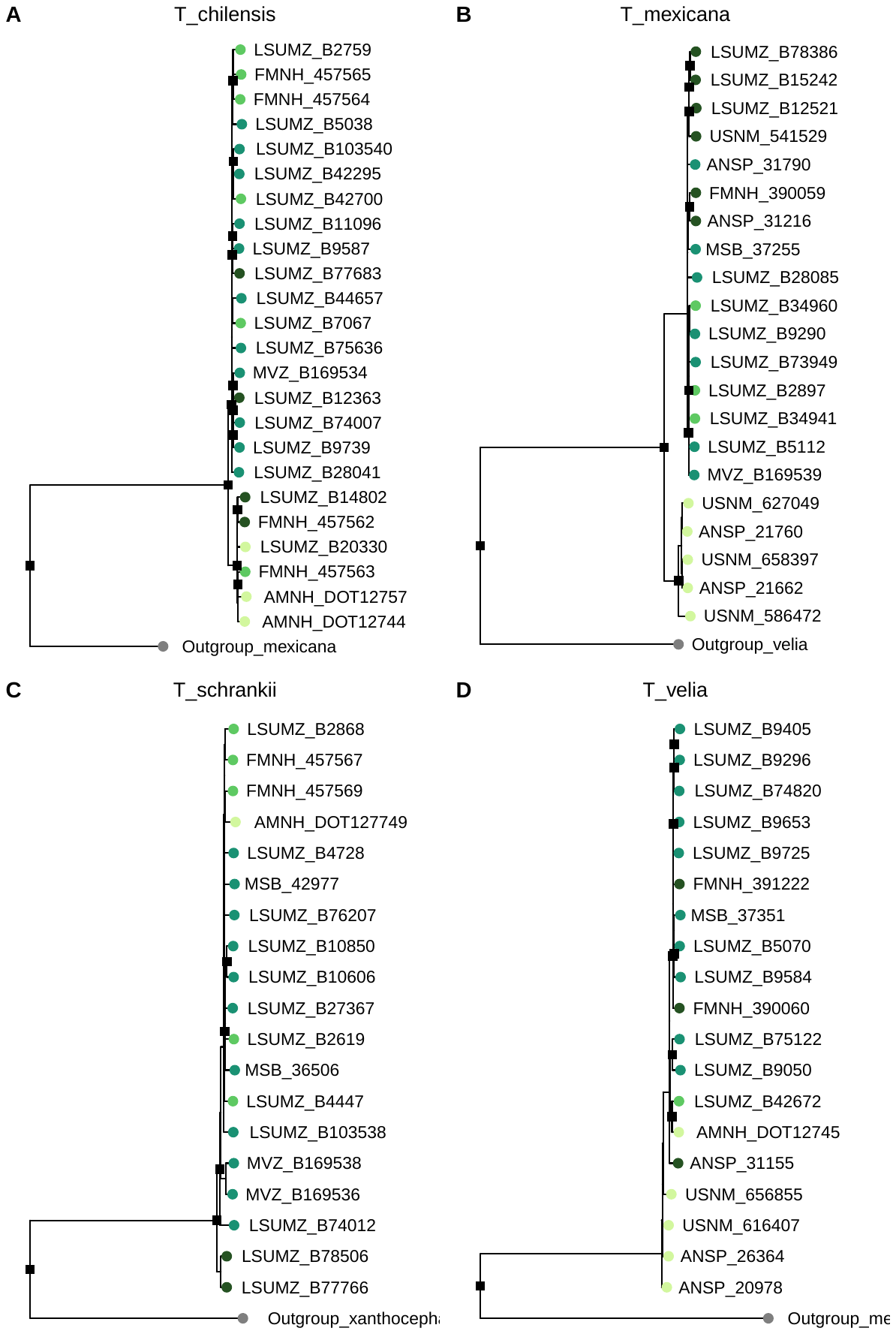

**Supplementary Figure S7.** Mitochondrial phylogenetic trees for the full 13 protein-coding gene sets for all Andean species. Trees were made with IQ-Tree 2 using automatic model selection from alignments partitioned by gene and codon, with 1000 ultrafast bootstrap replicates. Tips are color labeled according to their geographic population, as in Supplementary Figure 1. Nodes with > 80% bootstrap support are labeled with a solid black square.

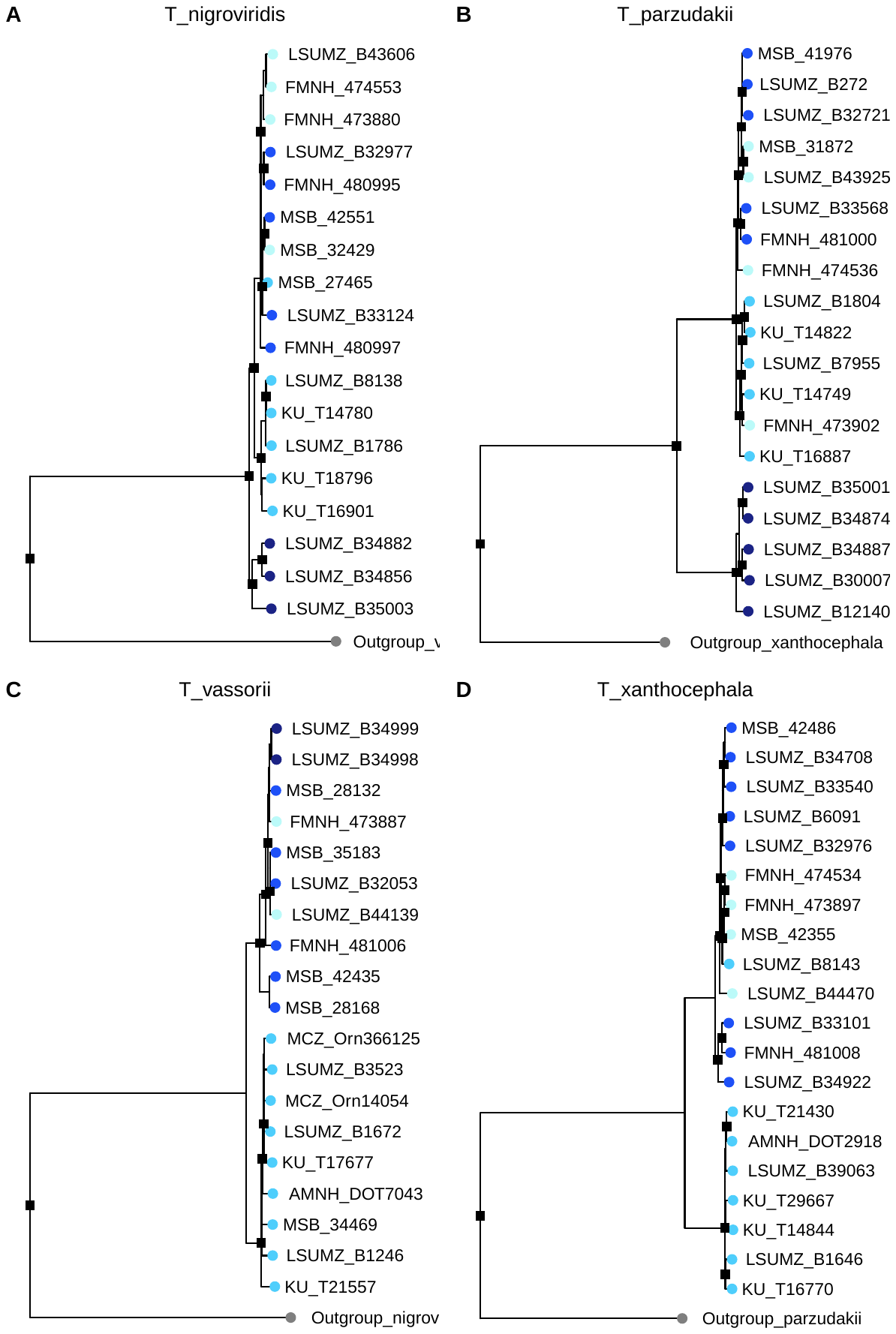

**Supplementary Figure S8.** Alternative geographic population designations for the Andes mountain system. A) Scenario 1 is the population system used in the main manuscript. B) Scenario 2 splits the “South Andes” population into two populations – “South Peru” and “South Apurimac”, separated by the Apurímac/upper Ucayali Valley barrier. The gray region is the dry western slope of the Andes that is not included in this study.

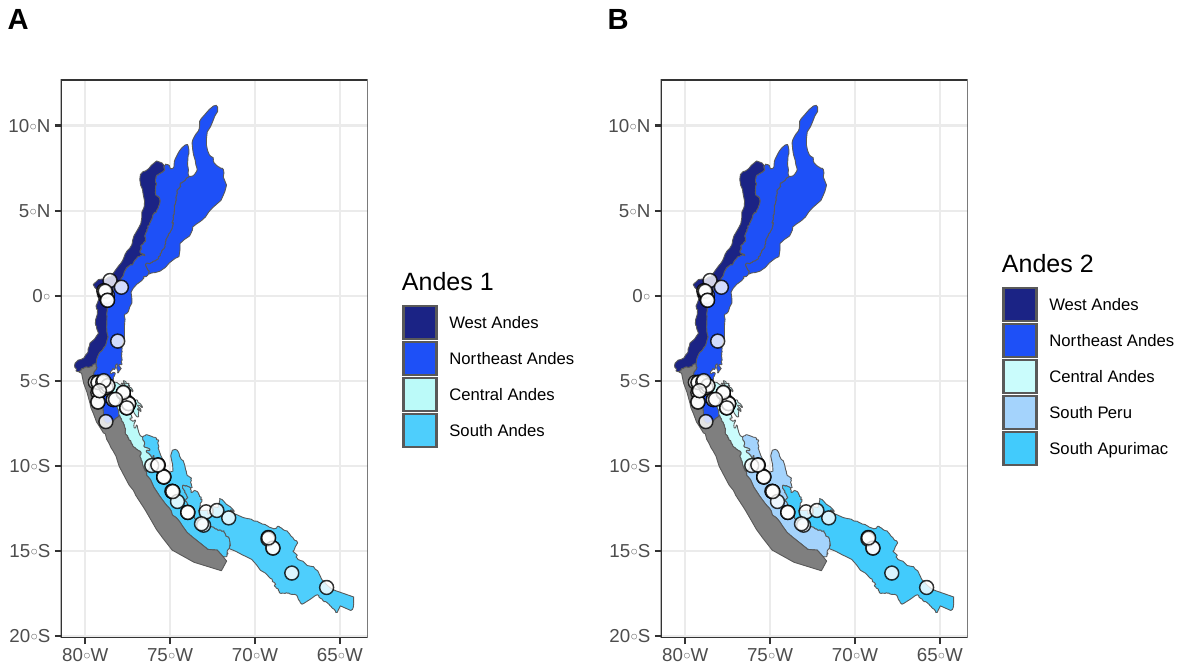

**Supplementary Figure S9.** Alternative geographic population designations for the Amazonian lowland system. A) Scenario 1 is the population system used in the main manuscript, with the Marañón River (the main source of the Amazon River) serving as the boundary between “North” and “Southwest” Amazon populations. B) Scenario 2 alternatively uses the Ucayali River, a wider headstream of the Amazon River, as the boundary separating “North” from “Southwest”. C) Scenario 3 isolates the region of western Amazonia between the Marañón and the Ucayali as its own geographic population, called “Huallaga.”

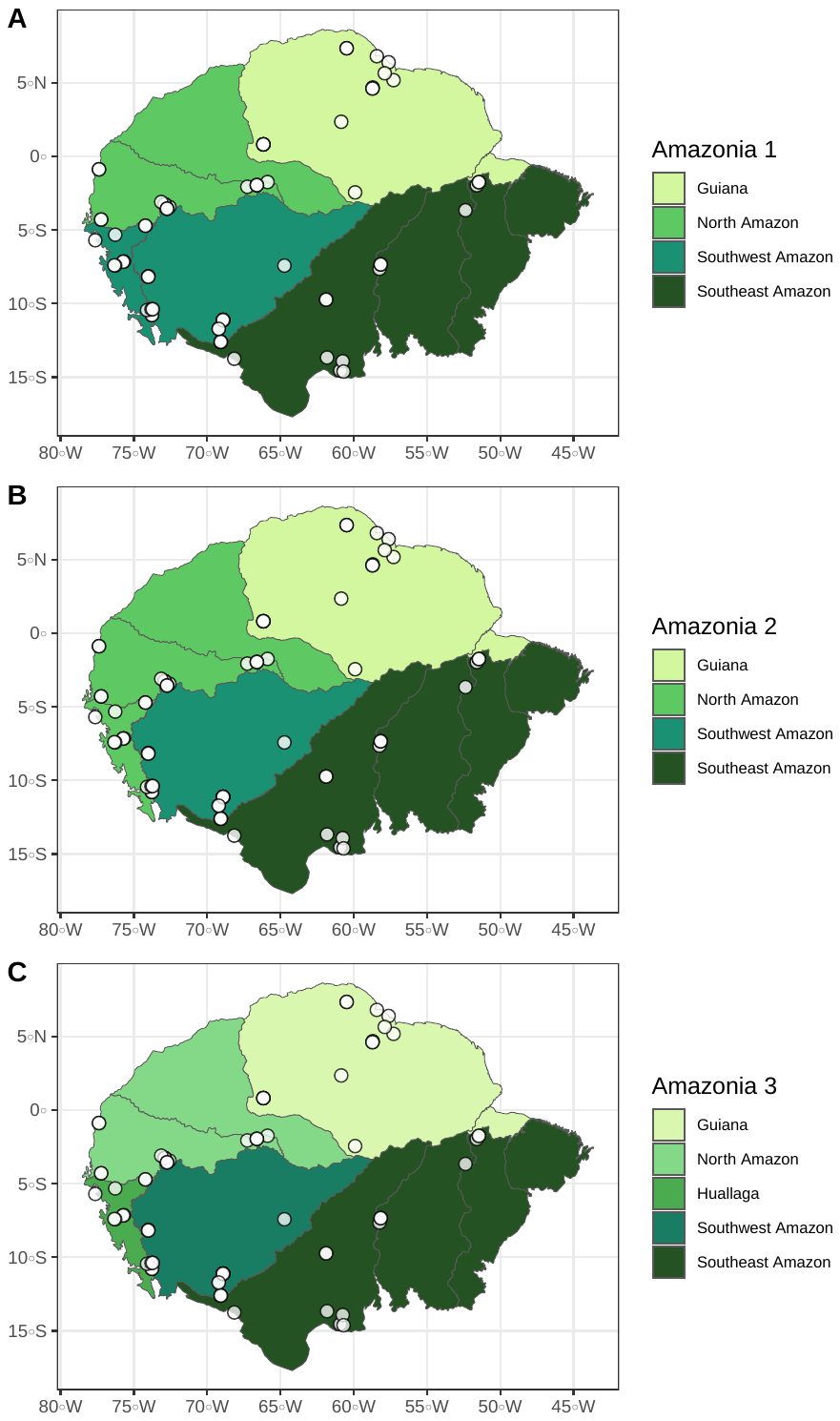

**Supplementary Figure S10.** Pairwise population divergence (F_ST_) across individual barriers, using alternative geographic population designations. There are two different Andes scenarios, and three different Amazonian scenarios, described in Supplementary Figures 8-9. For all 6 possible comparisons of alternative geographic scenarios, F_ST_ is significantly greater across barriers in the Andes mountains than barriers in the Amazonian lowlands.

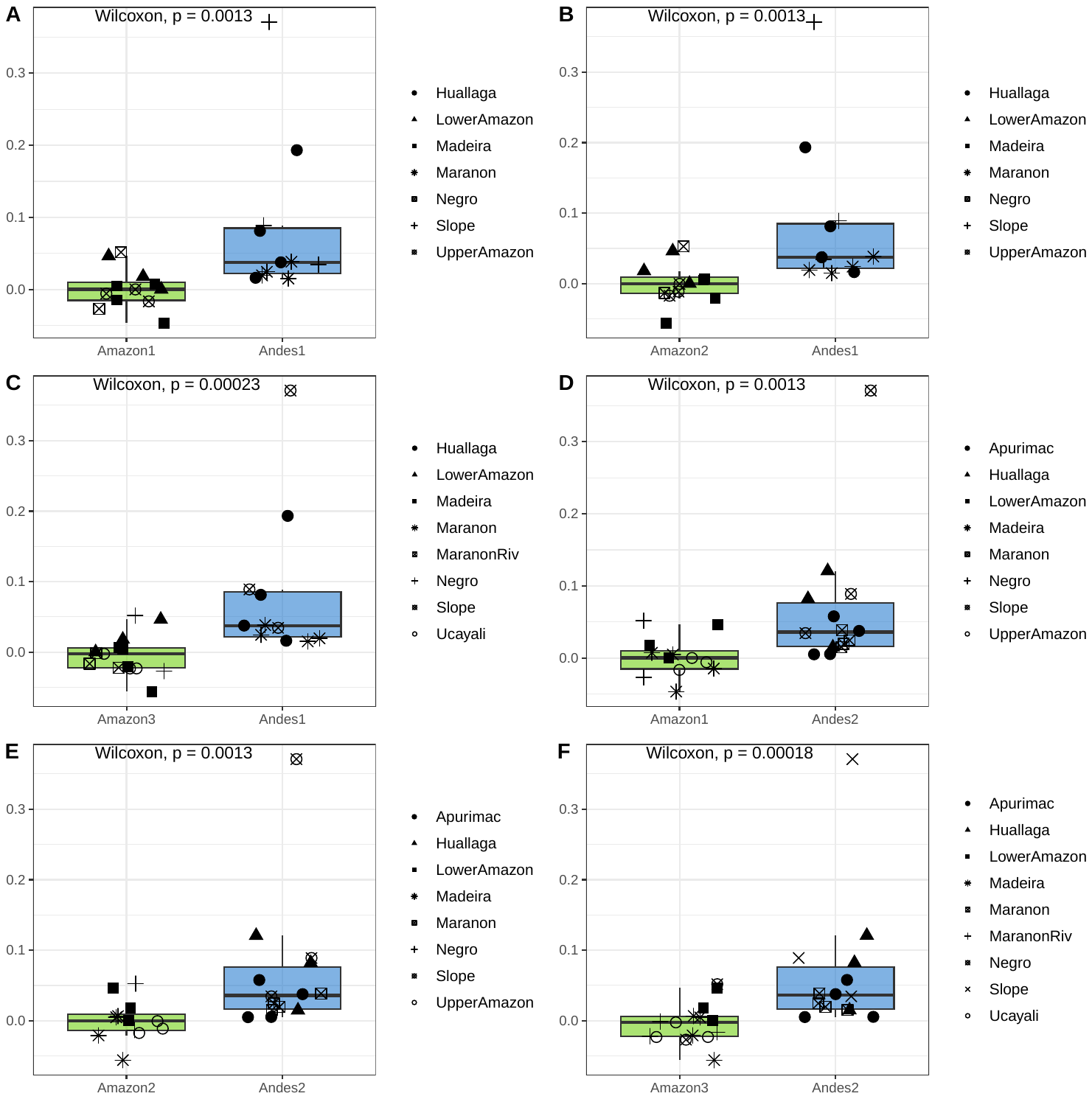

**Supplementary Figure S11.** Net migration rate (*m*) across individual barriers in the Andes mountains versus the Amazonian lowland. The average migration rate, measured as the percentage of the populations comprised of migrants per generation, is lower in the Andes than in the Amazon, but the difference is marginally non-significant.

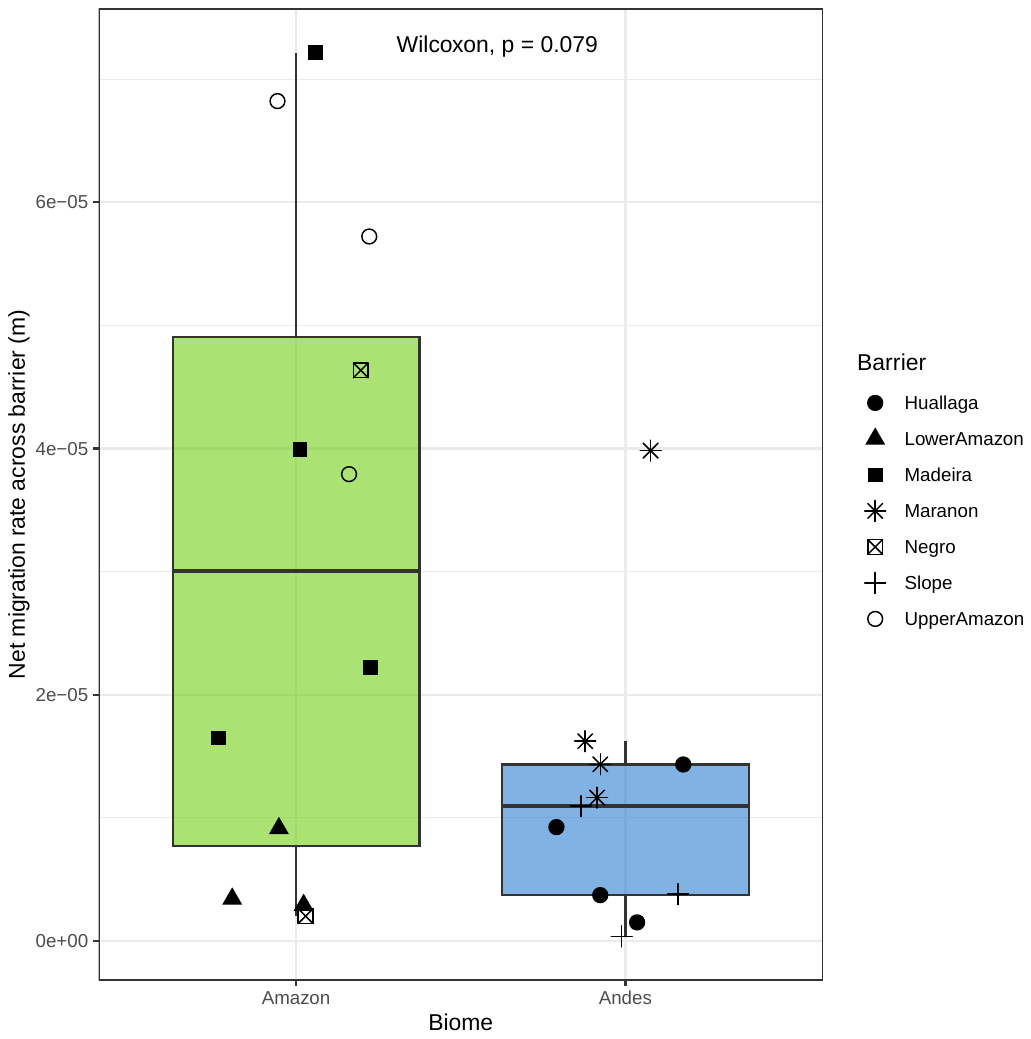

**Supplementary Figure S12.** An alternative metric of genetic diversity, pairwise nucleotide diversity (θ_π_), is also significantly greater in the Amazonian species than in Andean species (Wilcoxon test, p = 0.0286).

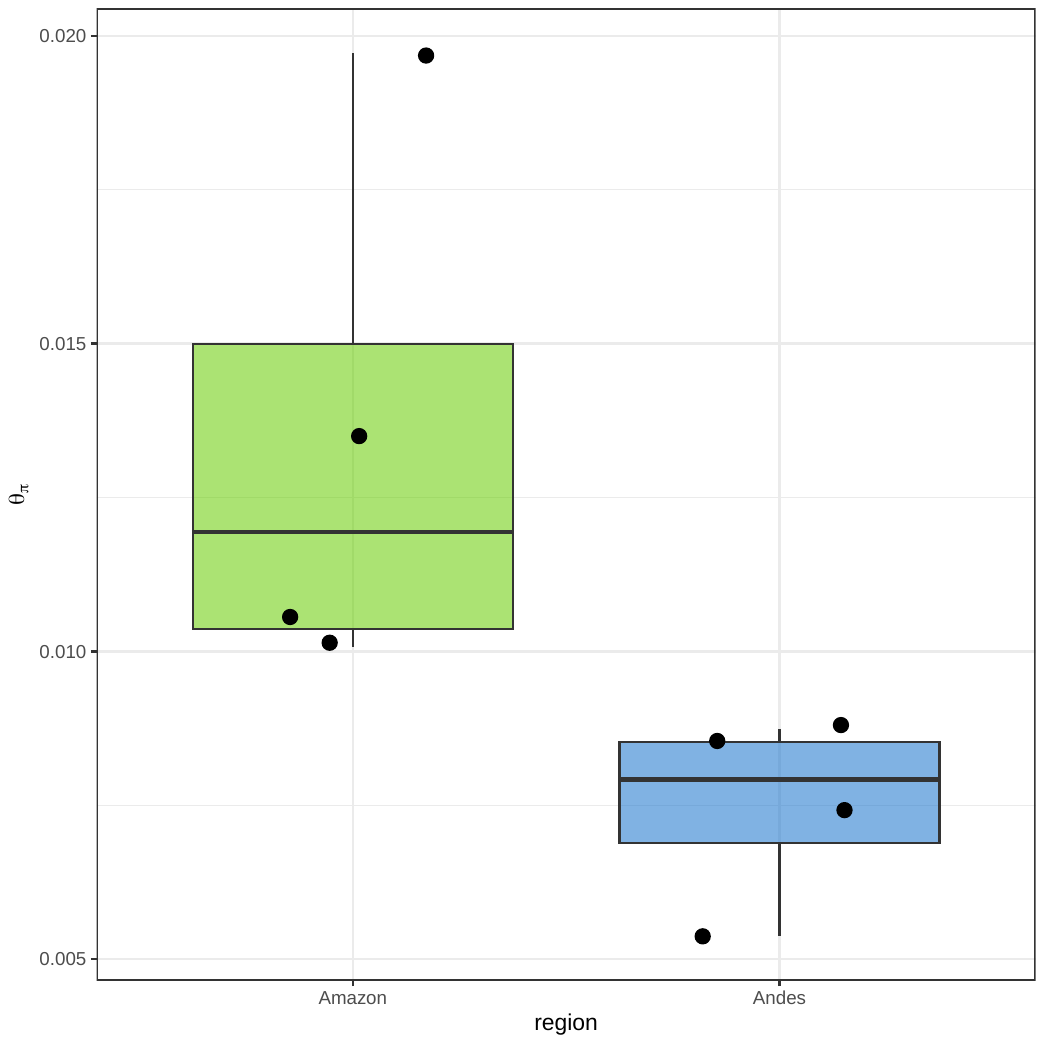
